# Large-scale Annotation of Biochemically Relevant Pockets and Tunnels in Cognate Enzyme-Ligand Complexes

**DOI:** 10.1101/2023.03.29.534735

**Authors:** O. Vavra, J. Tyzack, F. Haddadi, J. Stourac, J. Damborsky, S. Mazurenko, J. Thornton, D. Bednar

## Abstract

Tunnels in enzymes with buried active sites are key structural features allowing the entry of substrates and the release of products, thus contributing to the catalytic efficiency. Targeting the bottlenecks of protein tunnels is also a powerful protein engineering strategy. However, the identification of functional tunnels in multiple protein structures is a non-trivial task that can only be addressed computationally. We present a pipeline integrating automated structural analysis with an *in-house* machine-learning predictor for the annotation of protein pockets, followed by the calculation of the energetics of ligand transport via biochemically relevant tunnels. A thorough validation using eight distinct molecular systems revealed that CaverDock analysis of ligand un/binding is on par with time-consuming molecular dynamics simulations, but much faster. The optimized and validated pipeline was applied to annotate more than 17,000 cognate enzyme-ligand complexes. Analysis of ligand un/binding energetics indicates that the top priority tunnel has the most favourable energies in 75 % of cases. Moreover, energy profiles of cognate ligands revealed that a simple geometry analysis can correctly identify tunnel bottlenecks only in 50 % of cases. Our study provides essential information for the interpretation of results from tunnel calculation and energy profiling in mechanistic enzymology and protein engineering. We formulated several simple rules allowing identification of biochemically relevant tunnels based on the binding pockets, tunnel geometry, and ligand transport energy profiles.

## 1. Introduction

Enzymes are biological catalysts that can accelerate chemical reactions, which makes them essential for every living cell. These chemical reactions occur in the active site, which consists of residues with specific physicochemical properties. Active sites can be found either in clefts on the surface of an enzyme or buried inside a cavity shielded from the outer environment. In the latter case, the active site cavity is connected with the surface by access tunnels to enable the passage of ligands, small molecules that interact with the enzyme ^1^. This encompasses the exchange of reactant and product molecules or the binding of cofactors. The tunnels also impact the activity and specificity of the enzyme by restricting access to the active site for unfavourable molecules ^2^.

Several computational tools were developed for the detection of important cavities and pockets, e.g., FPOCKET ^3^, CASTp ^4^, and P2Rank ^5^. These tools rank all the pockets found in a protein structure by their scoring functions and select the best potential binding pocket for the user. To improve the reliability of the selection, one can use annotations found in structure databases ^6,7^. Unfortunately, these annotations are available only for a limited number of enzymes. The selection of the functionally relevant pocket is also crucial for the calculation of access tunnels. However, currently there is no tool available that would predict the suitability of a pocket for this purpose.

To identify tunnels in enzymes, one may use tools such as CAVER ^8^, MOLE ^9^ or MOLAXIS ^10^. Similarly, with pocket calculation, these tools can detect multiple tunnels and also provide ways to rank them based on their geometrical properties. In many proteins with buried active sites, multiple tunnels can be identified, which makes it difficult to decide which tunnel is biochemically relevant. This crucial decision could be greatly supported by a large-scale analysis of protein structures. Previous efforts in this matter focused purely on finding tunnels in enzymes ^11,12^. While these studies proved that tunnels are found in all enzyme classes, they did not define how to recognise biochemically relevant tunnels.

The classical computational approach to studying the biological relevance of tunnels is to simulate the interactions between a protein and a ligand with methods based on molecular dynamics ^13^. Unfortunately, this time-demanding type of simulation is not feasible for large datasets. More recent tools, such as CaverDock ^14^, GPathFinder ^15^, or ART-RRT ^16^, employ various approximations to simulate ligand transport in short computational times and provide valuable information about the energy profile of the process. These tools are gaining popularity ^17^ and have successfully been used for screening and identifying novel drugs ^18,19^ and protein engineering ^20–24^.

In this study, we present a novel strategy for annotating pocket relevance for tunnel calculation and assign biochemical relevance of tunnels based on ligand transport and binding energies. With the growing number of available protein structures ^25^ and models ^26^, automatic annotation of binding pockets and tunnels without the dependency on residue annotations would be of great use. Based on the premise that substrate and product molecules are present in relevant pockets in enzyme structures, we created a dataset independent of annotations. We selected experimentally derived enzyme structures with bound molecules that were similar to cognate ligands, i.e., ligands that potentially bind or react with a given enzyme. For this purpose, we used a previously published dataset of enzyme cognate ligand pairs ^27–29^, which we updated and utilized for structural analyses of pockets and tunnels in this study. We then developed a pipeline combining machine learning, the geometrical analysis of tunnels and the energy profiling of transported ligands. The pipeline was then validated against molecular dynamics simulations and applied to the large-scale dataset with more than 17,000 protein structures.

## 2. Methods

### 2.1. Automatic annotation

#### 2.1.1 Input data and annotation pipeline

The study uses data collected from the publication by Tyzack *et al.*^29^ from 2018 and updated in 2021 for the purpose of our study (Figure 1). The data consists of enzyme-ligand complexes ranked by the similarity of the bound ligand with the cognate ligand from the KEGG ^30^ database calculated by the PARITY algorithm ^29^. The following information is available for each case: KEGG Reaction number, EC Number, Protein PDB ID, Type of ligand (reactant or product), Bound ligand PDB ID, Cognate ligand KEGG number, and Similarity score (between bound and cognate ligands). In this analysis, we extracted all protein-ligand pairs with the Similarity score above 0.6, resulting in 35,882 entries. For annotation purposes, we only used unique PDB IDs, which pruned the dataset to 17,092 cases (Table 1).

**Figure 1:**
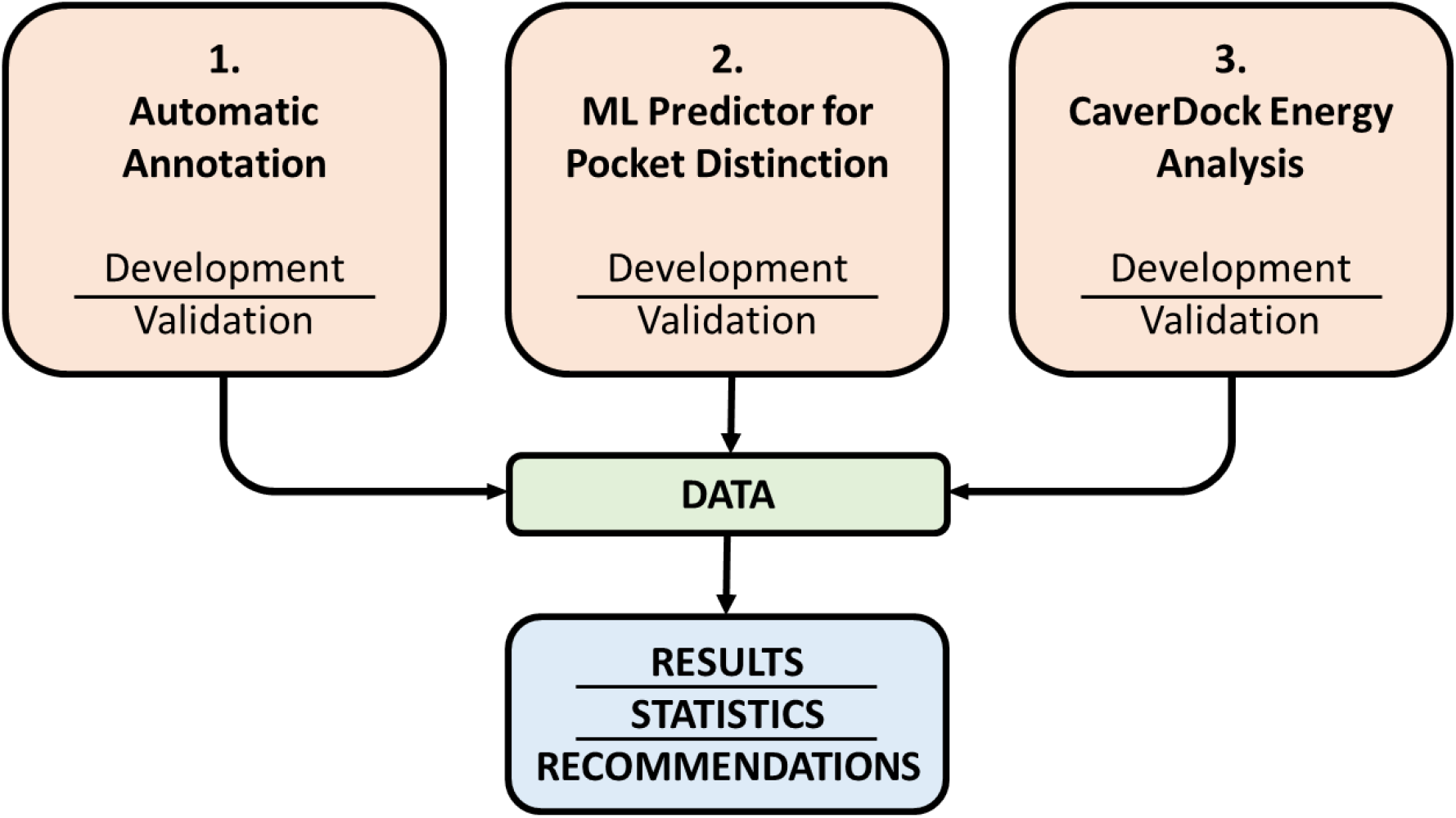
The overview of the pipeline developed in this study. The pipeline consists of three steps: (i) automatic annotation of enzyme-cognate ligand complexes with computational tools, (ii) classification of ligand binding pocket by ML predictor and (iii) analysis of ligand transport through enzyme tunnels with CaverDock.

**Table 1:** The summary of the pipeline proposed in this study and the dataset sizes at various stages of the pipeline execution.

| Pipeline | Items | Number of cases |
| --- | --- | --- |
| 1. Automatic annotation | Protein-ligand pairs | 35,882 |
|  | **Unique PDBs** | **17,092** |
|  | Ligand missing in the biological unit | 193 |
|  | Ligand not present in PDB | 133 |
|  | Ligand not present in any pocket | 1,058 |
|  | Pocket calculation errors | 337 |
|  | **Successfully calculated pockets** | **15,697** |
|  | Proteins with annotation in CSA and Uniprot | 11,046 |
|  | Proteins without annotation | 4,651 |
|  | Selected pockets with matching annotated residues | 8,350 |
|  | No matching residues in the selected pocket | 2,696 |
|  | No tunnels found by Caver | 526 |
|  | Tunnel calculation errors | 739 |
|  | **Successfully calculated tunnels** | **14,432** |
| 2. Machine Learning predictor for pocket annotation | Buried pockets without tunnels | 508 |
|  | Borderline pockets without tunnels | 160 |
|  | Surface pockets without tunnels | 597 |
|  | **Buried pockets with calculated tunnels** | **3,552** |
|  | **Borderline pockets with calculated tunnels** | **3,178** |
|  | **Surface pockets with calculated tunnels** | **7,702** |
| 3. Energy profiles | Protein-ligand pairs for CaverDock calculations | 14,432 |
|  | Unfinished CaverDock calculations | 1,274 |
|  | **Successfully completed CaverDock jobs** | **13,158** |
|  | Successfully calculated energy profiles | 29,693 |

The pipeline starts with running the annotation module of HotSpot Wizard ^31^ to search Swiss-Prot, UniProtKB ^6^, and CSA ^7^ for residue annotations using the sequence from the input PDB ID as a query. Moreover, the HotSpot Wizard module calculates solvent accessibility for each residue using the Shrake and Rupley algorithm ^32^ with BioJava ^33^. The first assembly containing the biological unit is downloaded from PDBe ^25^ in mmCIF format. The list of known cofactor three-letter codes is assembled based on the list of cofactors from FPOCKET ^3^ and CoFactor Database ^34^. Subsequently, the structure is cleared of any heteroatoms that do not belong to cofactors or bound ligands. The CIF file is then converted to PDB using the script cif2pdb from *gist.github.com/sbliven*. Each object from the original CIF is saved with a unique chain and matched by coordinates with the chains in the original PDB. From this point forward, only the PDB containing the biological unit is used in the rest of the pipeline, and only the first protein chain containing the bound ligand is analysed. When the biological unit is not available in PDBe, only the PDB within the asymmetric unit is used.

Next, pockets in the protein PDB are calculated using FPOCKET ^3^ with the following parameters: -m 2.8 (minimum radius an alpha sphere might have in a binding pocket) -n 10 (number of alpha spheres a pocket has to have close to alpha spheres of another pocket in order to be clustered together) -r 4.5 (parameter influencing the clustering of small pockets to larger pockets) -s 2.5 (parameter for multiple linkage clustering), and the list of the pockets related to the analysed protein chain is saved. In the next step, the information on whether these pockets contain the bound ligand is analysed with a simple in-house script. The script uses dummy spherical atoms from the files containing the 3D representation of each pocket. Then the atom coordinates of the bound ligand are checked to determine whether they are inside the dummy spheres. The information about how large a portion of the ligand is present in each pocket is collected and evaluated. In the situation where only one pocket is found to contain the ligand, this pocket is saved as the binding pocket. If multiple pockets contain the ligand (depending on the size and position of the ligand and pockets), the pocket with the largest portion of the ligand is selected if it contains at least 10 % more ligand atoms than the other pockets. In the case where there is less than a 10 % difference (for example, half of the ligand is in one, the other half in the second pocket), the pocket with a higher druggability score from FPOCKET is selected. Then the pocket residues are checked if they were annotated by the HotSpot Wizard module in the previous step. This step is done for validation purposes.

Finally, the selected pocket is used for setting up the starting point for tunnel calculation by CAVER ^8^. First, the geometrical centre of the protein biological unit is calculated, and based on the distance to Cα atoms of each pocket residue, the closest pocket residue is selected. Next, the geometrical centre of the pocket is calculated, and in the previously selected residue, the nearest side-chain atom closest to the pocket centre is found. After that, the coordinates of this atom are shifted by 0.5 Å in the direction of the vector connecting the atom and the centre of the pocket. After saving the coordinates for the tunnel starting point, CAVER is initialised with a probe radius of 0.9 Å and default remaining settings. When the tunnel calculation finishes, the results are collected and evaluated. The first five tunnels are saved based on the priority score from CAVER.

#### 2.1.2 Validation of Annotations

We validated the output from the pipeline in two ways. First, we analysed whether the residue annotations from UniProt and CSA were available for the selected binding pockets. We analysed the cases with available annotations from the databases and calculated the overlap to see how precise we were in selecting the correct binding pocket. Then we continued with the analysis of pockets categorised by the ligand coverage scenarios. We looked at how often the selected pocket had the best Fpocket and druggability scores in the matching/mismatching/no annotations subsets. This validation would show if the pipeline is universally usable for ligand-free structures.

Second, we studied the impact of the ligand present in the protein structure on the tunnel quality and geometry to evaluate the applicability of the pipeline for proteins without residue annotations or proteins without bound ligands. We used the REST API in PDBe ^25^ and RCSB ^35^ to search for identical protein structures not containing any bound ligands apart from water molecules, using 100% identity and the E-value cut-off of 1e^-15^. Only the PDB-IDs of proteins with successfully calculated tunnels during the annotations were used for the search. In the cases where we found multiple empty structures for matching enzyme-ligand complex from the dataset, we picked the first PDB in the list. We downloaded the biological units for the ligand-free structures and aligned them to the complexes with DeepAlign ^36^. Then we utilised the two structures as snapshots of one system and calculated the tunnels with the same settings as in the annotation pipeline. Using this workflow, we collected and analysed 2 904 pairs of protein-ligand complexes and ligand-free structures. To assess the structure pairs, we looked at the number of tunnels found in each pair and the tunnel properties provided by CAVER 3.02. We used the priority score from CAVER 3.02 that ranks the tunnels calculated by the tool. The priority score is calculated as a sum of the throughputs of all pathways in a given cluster, divided by the total number of snapshots that were analysed. In our case, the cluster of tunnels means that the same common tunnel was found in both structures in the pair. The throughput is calculated based on the cost function that takes into account the width and length of the tunnel, and it takes a value between 0 and 1, equal to the probability that the pathway is used as a route for the transportation of the substances ^8^.

### 2.2. Machine-learning predictor for pocket distinction

We initially used a simple metric of the ratio between buried and solvent-exposed residues in the binding pocket to assess its relevancy for tunnel calculation. However, preliminary data analysis showed that this approach was not sufficient. Since none of the available pocket calculation tools offers any metric describing the pocket being buried or exposed on the surface, we developed a new predictor to discriminate between the two.

We manually labelled 200 proteins with calculated pockets from the dataset for training. We analysed the distribution of EC classes in the dataset and randomly collected the number of proteins from each EC class following the distribution. The features for training the predictor were collected from the output of FPOCKET for the selected pockets, calculated in the annotation pipeline. These features represent the accessible surface area, polarity, volume, hydrophobicity, and other properties of pockets. Moreover, we kept the Exposed ratio as an extra feature. This ratio is equal to the number of exposed solvent-accessible residues as a fraction of the total number of pocket residues. The total number of features used was 20 (Table S1). We labelled each pocket by three classes: −1 for buried samples, 0 for borderline samples, and 1 for surface samples, according to manual visual inspection of pockets and protein structures. The borderline case was introduced primarily due to occasional difficulty assigning labels to some binding pockets. We also considered combining 0 and −1 classes into one to check if this would improve the results. We realise that manual annotation of accessibility is unreliable but it was impossible to generate the labels with other computational tools.

We used the Support Vector Machine (SVM), K-Nearest Neighbor (KNN), Shallow Neural Network (ANN), Gaussian Naive Bayes, and Random Forest classifiers. We applied GridSearchCV with 5-fold cross-validation for tuning hyperparameters of the algorithms from Scikit-learn: the function loops through predefined hyperparameters and fits the model to the training set to select the best parameters from a list based on the cross-validation error. The tuned hyperparameters and their corresponding ranges are listed in Table S2. For pre-processing data, we also tried a two-sample Kolmogorov-Smirnov test^37^. We used the Python function ks_2samp and the threshold of 0.11 to remove features scoring below the threshold from the dataset. Based on our results, we removed Apolar SASA, Volume, Alpha sphere density, and Alpha Sphere max dist for the two-class dataset. We tried three scenarios: (i) keep all three classes and features and tune the hyperparameters, (ii) absorb the borderline cases in the Buried class, remove the features based on the Kolmogorov–Smirnov test, and tune the hyperparameters, and (iii) the second scenario without the feature removal. We used accuracy, precision, recall, and F1 measure to report the performance of our predictor because the dataset was balanced:

Accuracy is defined as the percentage of correct predictions for the test data:

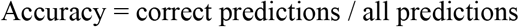

F1 measure is calculated from the precision and recall:

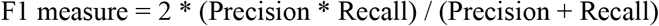

FPR is a false positive rate, which for a two-class problem is given as follows:

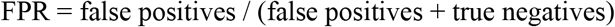

For the three-class problem, the most critical errors for our application are the cases when a pocket was assigned the label “1” (=surface) while the actual label is “-1” (=buried), as this will lead to falsely skipping the tunnel calculation step. Therefore, we used the following FPR formula for this case:

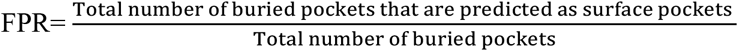

To establish the statistical confidence of our evaluations, we ran the code 100 times with different random seeds and reported the average and standard deviation from these runs. To validate the performance of the predictors, an independent test dataset with 100 samples was additionally collected and labelled in the same way as the training set (Table S3). The ratios of the samples in each class was similar to the train set. The selected best predictor was then used to classify all the calculated pockets.

### 2.3. CaverDock Energy Analysis

#### 2.3.1 Preparing CaverDock calculations

CaverDock was used to analyse the ligand pathways in all cases in the dataset with successfully calculated tunnels. For each calculation, the receptor, ligand, and tunnel input files were prepared in several steps: (i) The ligand was prepared by automatic parsing of the annotated cognate reaction and cognate ligand as follows. The annotated KEGG ^30^ reaction was used to build the reaction scheme. Then all of the reactants and product KEGG codes were parsed from the specific KEGG reaction website. Finally, using the molecule codes, the reaction scheme was written in a file (e.g., C01 C02 >> C03 C04). (ii) The molecule codes from the reaction scheme were used to parse the MOL file for each molecule. The MOL files were converted to SMILES codes, and the reaction was built based on the reaction scheme. (iii) The reaction in SMILES format was used as input in RD-toolkit ^38^. The RD-toolkit analysed the reaction and annotated the atoms which changed in the molecules. The algorithm in RD-toolkit can annotate reactions where the reactants and products differ in atom composition. It fails on isomerisation reactions because there is no change in the number or type of atoms. Moreover, the algorithm is unable to work with polymeric structures because of the formating in MOL files (such as cellulose). (iv) RDKit from *https://github.com/rdkit/rdkit* was used to decode the changes in the previously saved reaction and define the reaction centre for the molecules. (v) The information about the reaction centre was used for setting up the drag atom for CaverDock calculations. The drag atom is used to constrain the ligand to discs and pull the ligand through the tunnel. For reactants, the atom closest to the reaction centre was selected. In the case of products, the atom which was the most distant from the reaction centre was used. Pulling the ligand by the distant end of the molecule aims to better simulate the behaviour of leaving product molecule in the static snapshot of the protein structure. In the case of previous failure to annotate the reaction, the default settings were used (the atom closest to the molecule centroid). The explicitly set drag atom improved the success rate of CaverDock calculations with large ligands with many degrees of freedom. (vi) The cognate ligand MOL file was converted to PDBQT by using the prepare-ligand4 script from MGLtools ^39^. (vii) Each tunnel was discretized using the Discretizer tool from the CaverDock package with default settings. (viii) The PDB of the biological unit was converted to PDBQT using the prepare-receptor4 script from MGLtools ^39^. (ix) Finally, the grid box around the tunnel and the configuration file were prepared by the prepare-config script from the CaverDock package. Only the lower-bound trajectory was calculated and analysed.

Several important energy values were extracted from the energy profiles manually for the validation dataset and automatically in the annotation pipeline. EBound is the minimum energy in the binding site, and it was extracted from the first half of the energy profile. EMax is the maximum energy of the entire energy profile. For the collection of EMax values, we did not consider the peak energy at the beginning of the energy profiles, which is likely to be an artefact caused by pushing the ligand too close to the bottom of the tunnel. These values were extracted manually in the case of the MD validation and later based on our testing, we excluded 33 % of the profiles automatically in the data analysis part. ESurface was taken from the last tunnel disk at the surface of the protein. Since the binding energy of the fully unbound ligand in AutoDock Vina is 0, in the case where ESurface was higher than 0, we changed it to 0. This is linked to the fact that if we could extend the tunnel enough, we would always approach 0 kcal/mol once the ligand had no interactions. If the surface energy was below 0, we kept this value due to the possibility of favourable binding at the surface of the protein. The energy barriers were then calculated as E_a_ = E_Max_ – E_Bound_ for the products and E_a_ = E_Max_ – E_Surface_ for the reactants.

#### 2.3.2 Validation of CaverDock trajectories by molecular dynamics

The CaverDock tool has been tested extensively and used on various datasets in previous publications ^18,19^. However, validation of the quality of predicted trajectories from CaverDock has not been done by any method approaches based on Molecular Dynamics (MD). We explored several approaches for the simulation of ligand binding or unbinding, and in the end, we selected the Adaptive Steered Molecular Dynamics (ASMD) ^40^, because we were able to implement this method for our purpose. The ASMD method applies constant external force on two atoms in the simulated systems. This can be used to either push two atoms apart or pull them together to simulate unbinding or binding of ligands, respectively. To proceed with the validation, we selected eight cases that had at least two well-defined tunnels and the cognate product bound inside (Table 2). The selected proteins had 2-4 well defined tunnels. All the branches of the same tunnel apart from the highest–priority one were omitted. Out of the eight selected enzyme-cognate ligand systems, seven cases are not part of the final filtered input dataset used for the automatic annotation and data analysis due to the updates of the input dataset based on changes in structural databases used to gather the data. These changes have had no impact on the comparison of CaverDock with MD simulations.

**Table 2:** The comparison of Potential mean force profiles obtained from ASMD simulations and energy profiles from single structure or averaged CaverDock calculations over snapshots from MD simulations.

| Case | Enzyme | Ligand | Number of tunnels | Match with static CaverDock | Match with averaged CaverDock |
| --- | --- | --- | --- | --- | --- |
| System #1 (PDB ID 1OTW) | pyrroloquinoline-quinone synthase | pyrrolo-quinoline quinone | 3 | partial | partial |
| System #2 (PDB ID 2BFN) | haloalkane dehalogenase LinB | *trans*-3-Chloro-2-propene-1-ol | 3 | yes | yes |
| System #3 (PDB ID 2RFY) | cellobiohydrolase | cellobiose | 3 | no | yes |
| System #4 (PDB ID 2UWH) | cytochrome P450 BM3 | 11,14,15-trihydroxyicosatrienoic acid | 3 | partial | yes |
| System #5 (PDB ID 4E2Z) | C-3'-methyltransferase | Se-adenosyl-L-selenohomocysteine | 3 | yes | yes |
| System #6 (PDB ID 5EDT) | cytochrome P450 CYP121 | (4S)-4-(5,5-dimethylcyclohex-1-en-1-yl)cyclohex-1-ene-1-carboxylate | 4 | no | no |
| System #7 (PDB ID 3ORW) | phosphotriesterase | N-(6-aminohexanoyl)-6-aminohexanoate | 2 | yes | yes |
| System #8 (PDB ID 5U6M) | UDP-glucosyltransferase | uridine 5'-diphosphate | 3 | yes | yes |

To prepare the complexes for the unbinding simulations, we selected the lowest-energy binding pose from the CaverDock analysis of the first tunnel. We also checked that the binding pose with similar energy and position could be found in the CaverDock trajectories from other tunnels. After extracting this starting pose, we added hydrogens to the molecule by Open Babel ^41^. The RESP charges for ligands were calculated by Gaussian09_E.01 within R. E. D. Server ^42^. The correct bond types were assigned by Antechamber, and Parmchk2 was used to generate the FCRMOD files ^43^.

The protein molecule was processed by Pdb4amber ^43^, and the protonation state of the protein was calculated by the H++ server at pH 7 and 0.1 M salinity ^44^. The complex for MD simulation was then prepared with the following steps. The original crystallization solvent and the binding pose of the ligand were added, solvent molecules clashing with the position of the ligand were removed, and the *tLEaP* program of AmberTools 16 ^43^ was used to prepare the topology and coordinates files. The ff14SB force field ^45^ was applied in all simulations, Na^+^ and Cl^-^ ions were added to neutralize the system and achieve a 0.1 M concentration of NaCl, and an octagonal box of TIP3P ^46^ water molecules, with the edges at least 10 Å away from the protein atoms, were added.

The minimization and equilibration MDs were carried out with PMEMD.CUDA ^47,48^ module of AMBER 16 ^43^. In total, five minimization steps and twelve steps of equilibration dynamics were performed before the production of MDs. The first four minimization steps, composed of 2,500 cycles of steepest descent followed by 7 500 cycles of conjugate gradient, were performed as follows: (i) in the first one, all the atoms of the protein and ligand were restrained with a 500 kcal/mol·Å^2^ harmonic force constant; (ii) in the following ones, only the backbone atoms of the protein and heavy atoms of the ligand were restrained, respectively, with 500, 125, and 25 kcal/mol·Å^2^ force constants. A fifth minimization step, composed of 5,000 cycles of steepest descent and 15,000 cycles of conjugate gradient, was performed without any restraints. The subsequent MD simulations employed periodic boundary conditions, the particle mesh Ewald method for treatment of the long-range interactions beyond the 10 Å cut-off ^49^, the SHAKE algorithm ^50^ to constrain the bonds involving the hydrogen atoms, the Langevin thermostat with collision frequency 1.0 ps^-1^, and a time step of 2 fs. Equilibration dynamics were performed in twelve steps: (i) 20 ps of gradual heating from 0 K to 300 K, under constant volume, restraining the protein atoms and ligand with a 200 kcal/mol·Å^2^ harmonic force constant; (ii) ten MDs of 400 ps each, at constant pressure (1 bar) and constant temperature (300 K), with gradually decreasing the restraints on the backbone atoms of the protein and atoms of the ligand with harmonic force constants of 150, 100, 75, 50, 25, 15, 10, 5, 1, and 0.5 kcal/mol·Å^2^; (iii) 400 ps of unrestrained MD at the same conditions as the previous restrained MDs. The energy and coordinates were saved every 10 ps.

Before we started with the biased unbinding simulations, we ran classical MD simulations of *System #3* Cellobiohydrolase with Cellobiose and *System #4* Cytochrome P450 BM3 with 11,14,15-Trihydroxyicosatrienoic acid to showcase the need for steered MD and approximative methods for the study of ligand unbinding. We used the prepared complexes and ran 3 replicas of 1 μs simulations to study the behaviour of the complexes and potential unbinding of the ligand molecules.

Next, the unbinding trajectories were calculated with ASMD ^40^. We used the settings obtained from the tutorial for AMBER. The following parameters were used: 25 parallel simulations, 2 Å stages, a velocity of 10 Å/ns, and a force of 7.2 N. The rest of the MD settings were set as in the last equilibration step. The protein atom for the steering was different for each tunnel. We selected the Cα atom in the residue at the bottom of the tunnel which was located opposite to the tunnel opening so that the ligand could be pushed from the binding site in this direction (Table S4). The ligand atom for steering was selected as the one closest to the centroid of the molecule. This setting was used in all systems, except for the *System #8* with UDP-glucosyltransferase and Uridine 5’-diphosphate: Due to the size and shape of this ligand, we were unable to simulate any trajectories using the centroid atom. In this case, we had to select the atom closest to the tunnel for each unbinding simulation. The selenium atom in Se-Adenosyl-L-selenohomocysteine of *System #5* was changed to sulphur in ASMD simulations. To compare the ASMD simulations with CaverDock, we extended the CaverDock trajectories to match the length of the simulated distances from ASMD.

Apart from the ASMD simulations, we ran MD simulations with ligand-free structures to generate ensembles of protein snapshots to study how much CaverDock results change when using dynamical structures. We used the same settings for the preparation of the systems, minimisation, and equilibration. We ran 50 ns of production MD, saved every 1,000^th^ snapshot, and collected 100 snapshots covering the entire MD simulation (every 250^th^). The solvent molecules were deleted, and the snapshots were aligned and centred to the first snapshot with Cpptraj ^43^. We calculated the tunnels in selected snapshots using CAVER 3.02 ^8^ with a smaller probe radius of 0.5 Å. The rest of the setup was the same as during the annotation. Tunnel clusters found in MD snapshots were visually matched with the tunnels found in the static structures. Using the same workflow for CaverDock simulations as in the annotation, we calculated the transport of ligands through the snapshots. The tunnels were discretized and extended by 5 Å. CaverDock calculations were run with the drag atoms defined in the annotation pipeline. Finally, we averaged the energy values for each tunnel in every system.

## 3. Results

### 3.1 Automatic annotation

#### 3.1.1 Annotation of the filtered PROCOGNATE dataset

The summary of the filtering of the PROCOGNATE dataset is given in Table 1. Out of the 17,092 unique PDBs, the ligand was not present in the biological unit in 193 cases, so we had to use the asymmetric unit instead. In 133 cases, the ligand was not present in the PDB at all – when three-letter codes for ligands did not match the bound ligand code in the dataset. In 1,058 cases, the ligand was not inside any of the calculated pockets but rather at or near the protein surface. In 337 cases, there were errors in the pocket calculation. We looked at the EC class representation in the 15,697 cases with successfully calculated pockets, and all EC classes were represented in the dataset: EC 1 (25.1 %), EC 2 (38.5 %), EC 3 (20.4 %), EC 4 (7.7 %), EC 5 (4.3 %), EC 6 (3.6 %) and EC 7 (0.4 %). Concerning the tunnel detection, no tunnels were found in 526 cases, and 739 cases finished with errors. This could stem from the following: (i) the pocket was at the surface of the protein and the CAVER algorithm was unable to calculate any tunnels, (ii) the automatically set starting point was in an incorrect position, or (iii) the space was too narrow for the 0.9 Å probe during the calculation, and the tunnel calculation failed.

#### 3.1.2 Validation of Annotations

The two-fold validation was used to evaluate the usability of the proposed pipeline. We analysed the collected annotations for residues essential for function, i.e., catalytic or binding residues, in selected binding pockets. By searching UniProt and M-CSA, we managed to find annotations for 11,046 protein structures, and for 4,651 structures, we found no information on essential residues (Table 1). Out of 11,046 annotated cases, 76 % matched the essential residues with the pocket-lining residues.

We further investigated the impact of the selection of the studied pocket on the performance of the pipeline. Using the ligand coverage, i.e., the fraction of the molecule overlapping with a pocket, we discriminated between three scenarios: i) the ligand belonged only in one pocket (single pocket), ii) a part of the ligand was found in another pocket, but the ligand was occupying the main pocket by 10 % more than other pockets, iii) the ligand occupied multiple pockets, and the difference was less than 10 %. In the third scenario, e.g., when half of a ligand was inside one pocket, and the second part lay in another (Figure S1), we selected the pocket with the highest druggability score. To this end, we looked at how often the selected pocket has the best Fpocket and druggability scores in the matching/mismatching/no annotations subsets (Table S5). In these subsets, the selected ligand-binding pocket was top-ranked by FPOCKET scores only in 43 %, 27 %, and 41 % of the cases. In the case of druggability scores, it was 23 %, 12 %, and 17 %, respectively. These values were surprisingly low, implying that selecting the pocket based on calculated scores would lead to a high number of errors. On the other hand, based on the 75 % overlap of the selected pockets with annotated essential residues in structural databases, we can say that the approach of selecting the pocket based on the ligand location is significantly better than a blind selection of the best pockets ranked by Fpocket score or druggability. Furthermore, using the same settings for the FPOCKET calculation for all proteins in the dataset seems insufficient as it led to cases where the ligand overlapped with multiple pockets. In addition, selecting the pocket by predicted scores is not generally applicable to any ligand-free structure without available essential residue annotations. A solution could be to extrapolate the location of the ligand and selected pocket from structurally similar proteins.

In the second part of the validation, we analysed how the presence of a ligand impacted the geometry of tunnels in proteins in pairs of ligand-bound and ligand-free structures. We used the priority score in CAVER 3.02 to calculate how many out of the top 5 tunnels identified in ligand-bound structures could also be found in the top 5 tunnels of ligand-free structures (Figure 2A). In the 2,904 studied pairs, we found no common tunnels in 24 % of the cases. This could be caused by the absence of the ligand in the structure, which led to a narrower binding site and impacted the geometry of calculated tunnels. In this category, no tunnels were found in the ligand-free structure in 146 cases, and only one tunnel, which did not match with any of the tunnels from ligand-bound structures, was found in 139 cases. In the rest of the structures, 35 % had one common tunnel. Based on the results, we observed that it was generally rare for a protein to have more than three potentially biologically relevant tunnels. We collected the priority scores for each of the five ranks of common tunnels and calculated the probability distribution to further study the clusters and define a metric for potentially relevant tunnels (Figure 2B). We concluded that the tunnels with the priority above the average of the third tunnel around 0.55 could be potentially relevant, with geometrical properties suitable for ligand un/binding. We suggest that for screening purposes, users should focus only on the first three tunnels calculated by CAVER 3.02 or use more tunnels with a priority score above 0.55. This recommendation is aimed only at the cases in which there is no previous information about the relevancy of tunnels in the studied protein. Based on these findings, we focused only on the first three tunnels in our subsequent data analyses.

**Figure 2:**
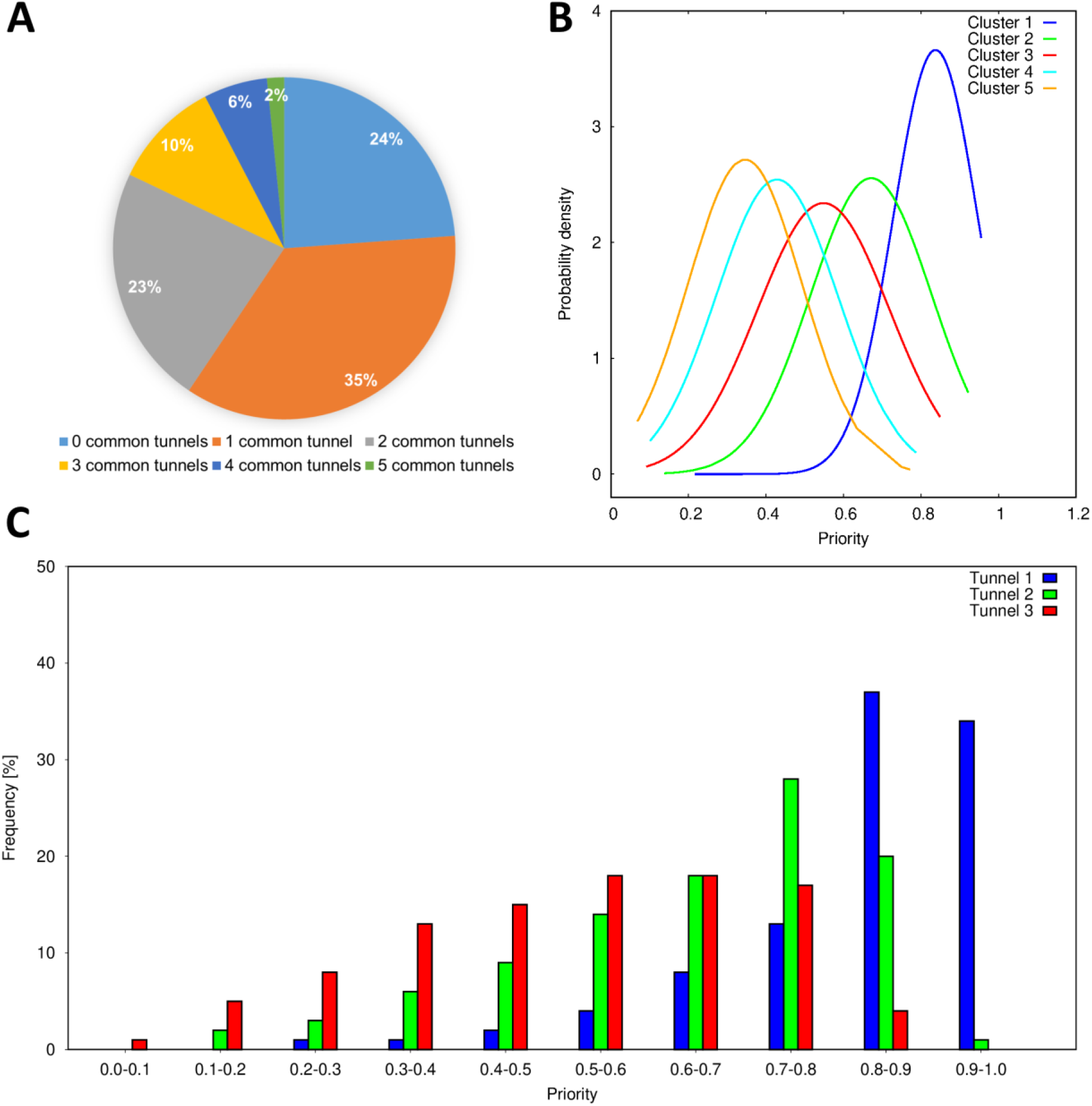
Analysis of tunnels in pairs of ligand-bound and ligand-free structures and the entire annotated dataset. (A) The number of common tunnels found in both ligand-bound and ligand-free structures. (B) Probability of distribution of CAVER priority score for the best five clusters in pairs of structures. (C) Distribution of the priority score for the first three tunnels in all proteins from the dataset with calculated tunnels. The analyses show that the first three tunnels are commonly present in enzyme-ligand complexes and ligand-free structures. These tunnels have the best geometrical parameters and are suitable for ligand un/binding. Tunnels with the priority above 0.55 could be potentially biologically relevant.

### 3.2 Machine-learning predictor for pocket discrimination

Since tunnel calculations are of little use for the surface binding pockets in the annotation pipeline, we trained a machine-learning predictor for their identification. We used KNN, Random Forest, SVM, ANN, and Naïve Bayesian to discriminate between buried and surface binding pockets. We tested two annotation strategies: a three-class problem (buried, borderline, surface) and a two class-problem in which the buried and borderline classes were merged into one (Table S6). Despite featuring lower absolute values, the three-class prediction results were similar to those for a two-class predictor if the baseline accuracy of a completely random prediction was taken into account (33% vs. 50%). The highest values of 1-FPR for both setups were close to each other: 75% in the case of KNN for the three-class predictor and 70% in the case of the ANN for the two-class predictor (SI-Table 4). We also checked if the misclassified cases of KNN and Shallow NN in the three-class dataset are the same, and they did not overlap except in a few cases, i.e. 8 buried pockets misclassified as surface in the test set. Moreover, while both predictors showed low performance in borderline cases and similar performance in the buried cases, the ANN predicted most surface pockets correctly. The Naïve Bayesian predictor was added as a simple baseline, and while it showed the highest value of 1-FPR of 90% and 93% on the training dataset for three- and two-class problems, respectively, it failed to identify any buried samples in the test dataset. In addition to the evaluation of our predictors on the test set, we built learning curves to decide whether expanding the training set might help improve the predictors (Figure S2). However, based on the curves, the accuracy was likely to stay around 50% even if more data points were added for training. Moreover, the feature pre-selection based on the two-sample Kolmogorov-Smirnov test did not improve the results, so we used the entire set of features in our final predictor. Based on the performance on the test set, we selected the ANN three-class predictor to annotate the successfully calculated pockets (Table 1).

### 3.3 CaverDock Energy Analysis

#### 3.3.1 CaverDock annotation results

We analysed 14,432 proteins with calculated tunnels with CaverDock. We were not able to produce ligand trajectories for 1,244 protein-ligand systems (Table 1) due to several factors: (i) we had problems with the automatic parsing of ligand data from KEGG, (ii) protein structures contained parts of DNA or RNA which caused the receptor preparation to fail, (iii) we failed to discretize the tunnels for CaverDock because they were extremely short, represented by only one dummy sphere or one sphere encompassed by another, or (iv) we discarded the cases in which the lower-bound CaverDock calculation did not finish within 48 hours on 4 CPUs. Based on the tunnel priority distribution, we analysed the energies of ligand un/binding in up to three tunnels found in each protein. In 13,188 successfully calculated protein-ligand systems, we produced 29,752 energy profiles: 12,804 trajectories for the tunnel 1, 9,465 for the tunnel 2, and 7,483 for the tunnel 3.

#### 3.3.2 MD simulations for validation of CaverDock trajectories

Both unbiased and biased MD simulations were used to validate the quality of CaverDock results. We simulated three replicas of 1 μs unbiased MD simulations for cellobiohydrolase with cellobiose and cytochrome P450 BM3 with 11,14,15-trihydroxyicosatrienoic acid (*System #3* and *#4* in Table 2, respectively). The ligand remained in the binding site, and we did not observe unbinding in any replicas. This result showed the importance of applying bias in MD to study events such as ligand unbinding. Furthermore, it demonstrated the applicability of approximative methods for the simulation of un/binding to save computational time and effort since unbinding was not observed even in these long simulations. We qualitatively compared the match between the Potential mean force profiles from ASMD and CaverDock calculations. We used either CaverDock trajectory from a single static structure or averaged CaverDock results from 50 ns MD snapshots (Table 2). We show the highest energy value in the profile EMax for the static and the averaged CaverDock calculations in Table S7.

In the case of *System #1* (Figure S3), the energies for tunnels 1 and 2 were similar, but the order was swapped compared to the ASMD simulations. Both tunnels were not frequently open in the 100 snapshots. Moreover, the priority of tunnel 1 was lowered in MD snapshots, so both tunnels 1 and 2 seem to be feasible for ligand binding. The ligand was not able to unbind through tunnel 3 in ASMD simulations, which agrees with the large barrier found in CaverDock energy profiles. Although the match remained partial even after analysing the averaged energies from MD snapshots, the EMax values were improved. Therefore, we concluded that this case was impacted by the rigidity of the crystal structure. *System #2* had perfect matches for all three tunnels (Figure 3). In ASMD the ligand was able to unbind with difficulties in tunnel 3, but the force started to unfold the part of the protein that was used for steering the simulation. This result is in accord with the large CaverDock barriers. In *System #3*, there was no match between ASMD and CaverDock results for the static structure (Figure S4). The use of averaged results from MD snapshots improved the matches. We concluded that the loops around tunnel 2 made it too wide open in the static structure and biased the results. The ligand in *System #4* unbound successfully in both tunnels 1 and 2 (Figure S5). On the other hand, it did not unbind through tunnel 3 and remained stuck in the binding site. There was only a partial match in the static structure but good agreement in averaged profiles. Therefore, we deduced that both tunnels could be preferred by the ligand. In *System #5*, there was no unbinding observed in tunnels 2 and 3 (Figure S6). The results from all the simulations agreed. The inability to pass through the tunnels in ASMD was reflected in the barriers in both types of simulations. *System #6* had no matches between CaverDock and ASMD, and the use of averaged energies from snapshots did not improve matches or EMax values (Figure S7). The crystal structure seemed too compact and did not have enough time to open during the short MD simulation of the complex. In *System #7*, the ligand was able to unbind successfully through tunnel 1 but was not able to pass through tunnel 2 (Figure S8). CaverDock results agreed with ASMD, so we had a good match across all simulations. In the case of *System #8*, the ligand preferred tunnel 1 over tunnel 2 and was not able to pass through tunnel 3 (Figure S9). Static CaverDock showed similar energies for both tunnels 1 and 2, and the results were improved in MD snapshots, where we saw a slightly higher barrier in tunnel 2. It indicated that both tunnels 1 and 2 could be used by the ligand.

**Figure 3:**
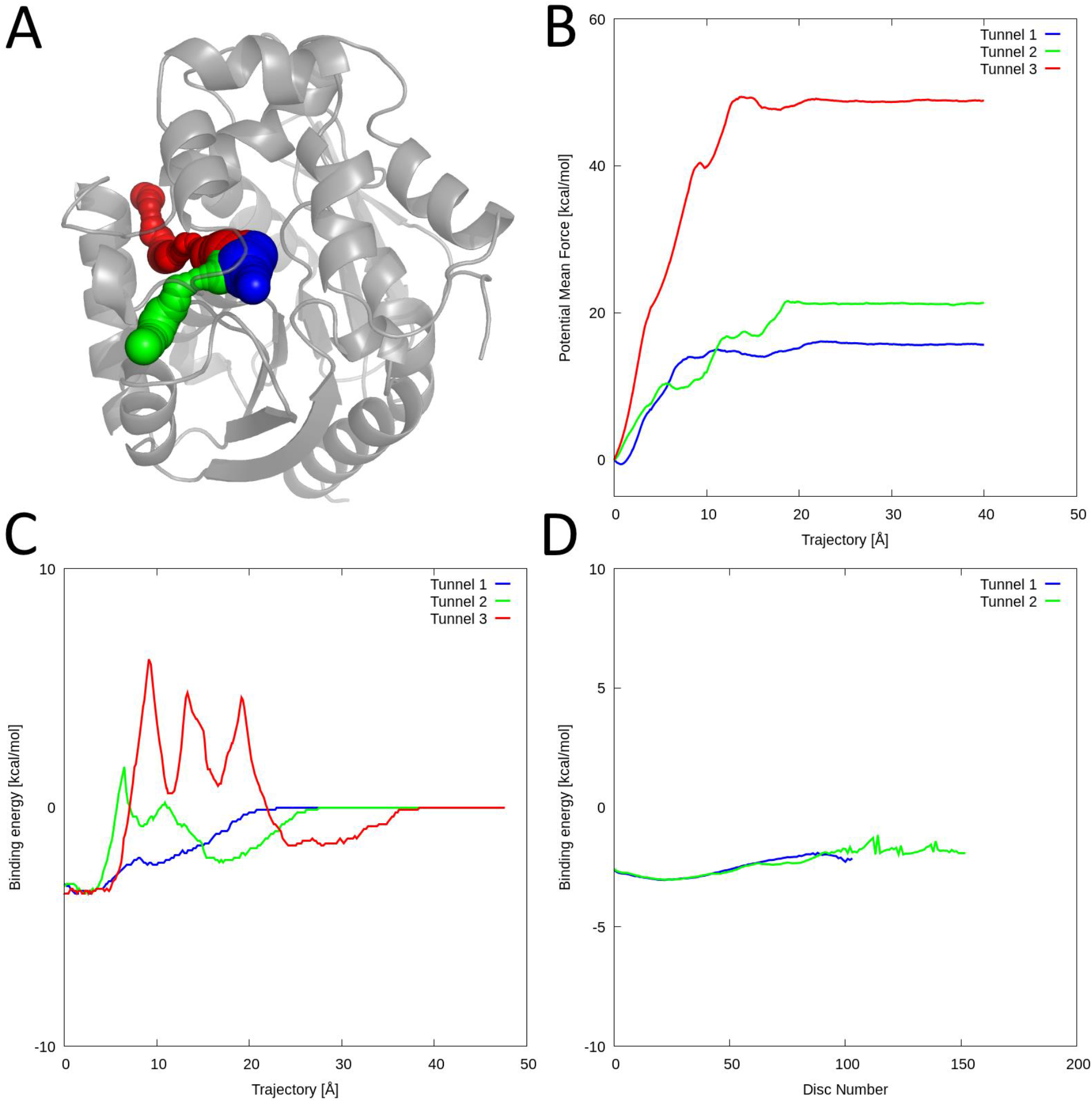
Results from CaverDock validation for haloalkane dehalogenase LinB with trans-3-chloro-2-propene-1-ol. (A) Visualisation of the protein structure (PDB ID 2BFN) with analysed tunnels showed as spheres: tunnel 1 (blue), tunnel 2 (green), tunnel 3 (red). (B) Potential mean force profiles from ASMD simulations. (C) Energy profiles from static CaverDock calculations. (D) Averaged CaverDock energy profiles from 50 ns simulation snapshots. The third tunnel was not present in the MD snapshots. The System #2 showed qualitative aggrement between the ASMD and CaverDock results.

### 3.4 Data Analysis

The ANN predictor was used to discriminate the pockets based on their type for all cases within the dataset. In the case of the pockets for which we did not manage to calculate tunnels, 508 pockets were predicted as buried, 160 as borderline, and 597 as surface. This was a surprising finding since we expected all these pockets to be predicted as surface pockets. In the second part of the dataset, i.e., the cases with pockets and successfully calculated tunnels, 3,552 cases were predicted as buried pockets, 3,178 as borderline, and 7,702 as surface (Table 1). In the subset of proteins for which we were able to identify pockets and tunnels, we coupled the predictions with the information about tunnels (Table S8). We binned tunnels similarly as in the study of Pravda *et al.* ^51^: short tunnels under 5 Å, medium-length tunnels between 5 Å and 15 Å, and long tunnels over 15 Å. In tunnel 1, we see a significant overlap between the categories of pockets with corresponding tunnel lengths. The vast majority of tunnels (75 %) in buried cases were either medium or long. Borderline cases were defined as a separate category because during manual annotation, it was difficult to assess if pockets were completely open on the surface or partially buried. For this category, we had 41 % short, 49 % medium, and 10 % long tunnels. For the surface cases, 74 % were short tunnels. Thus, our predictor proved successful in its predictions for tunnel 1 and could be a useful tool for assessing whether the calculation of tunnels in a protein makes sense or there is just a surface cavity. We carried out a similar analysis for tunnels 2 and 3, but these tunnels were of lower priority and always longer than tunnel 1. Therefore, almost all the tunnels were either medium or long. Based on this result, we defined tunnel 1 as the only reliable descriptor of the relationship between the predicted pocket type and tunnel length. Moreover, the proteins with tunnels shorter than 5 Å could potentially be discarded since they were calculated for pockets predicted as surface pockets and were, therefore, irrelevant to the tunnel analysis. The main benefit of the predictor is the possibility of pocket annotation in enzyme structures with very narrow tunnels, which would not be found unless the user used a smaller probe during the calculation, or when the tunnel calculation fails. One could also use the predictions to decide whether calculating and analysing tunnels is worthwhile for a particular protein structure. Since the predictor does not require the presence of a ligand in the structure, it is also generally applicable for ligand-free structures.

We studied the geometry of the first three tunnels in more detail. The distribution of tunnel priority scores for all cases with calculated tunnels is presented in Figure 1C. Importantly, we observed the same trend in the priority scores as in the analysis of pairs of complex and ligand-free structures. The throughput of tunnels 2 and 3 was lower because they were narrower, longer, and more curved than the tunnel 1 (Figure S10). This is not surprising since the priority score is related to the geometrical tunnel properties. Therefore, the priority score should be a sufficient metric for screening purposes. We continued this analysis by separating the dataset based on EC numbers (Figure S11). Tunnels were present in proteins from all EC classes, which was in agreement with previous studies ^11^. The tunnel priority followed the same trend in all the classes apart from EC 7 due to the low number of cases in the dataset. We did not observe any major differences in the geometrical properties, which would otherwise indicate that certain EC classes preferred tunnels with specific geometries. We also studied the number of tunnels in each EC class with a priority higher than 0.55 (defined in the analysis of pairs of structures). Apart from EC 7, the results were similar for all EC classes (Figure S12). For future tunnel analyses, it might be worthwhile to compare subclasses to see more significant differences in tunnel geometries.

Next, we studied whether the geometrical bottleneck, i.e., the narrowest part of a tunnel, was the best hot spot for mutagenesis to improve ligand binding and selectivity. For this purpose, we collected the maximum energy EMax from each CaverDock trajectory. In the next step, we compared the location of the energy maximum and the geometrical bottleneck in the tunnel (Figure 4). We tracked how often the maximum energy was in the disc with the lowest radius or in its vicinity (1.5 Å, 3 Å, and 5 Å). The match between the energy and geometry bottleneck was around 50 % for the exact disc and 75 % for the 5 Å vicinity (Table S9). The mismatch proved that the tunnel geometry is not a sufficient indicator alone to identify the bottleneck for the transport of a particular ligand. Analysis of the energy profile by approximative methods could help with the identification of other important hot spots for modification of ligand transport. The analysis was run with cognate ligands; therefore, these molecules should be recognizable by the enzymes. The results might change for a set of ligands of a larger size or with physicochemical properties different from the cognate ligands.

**Figure 4:**
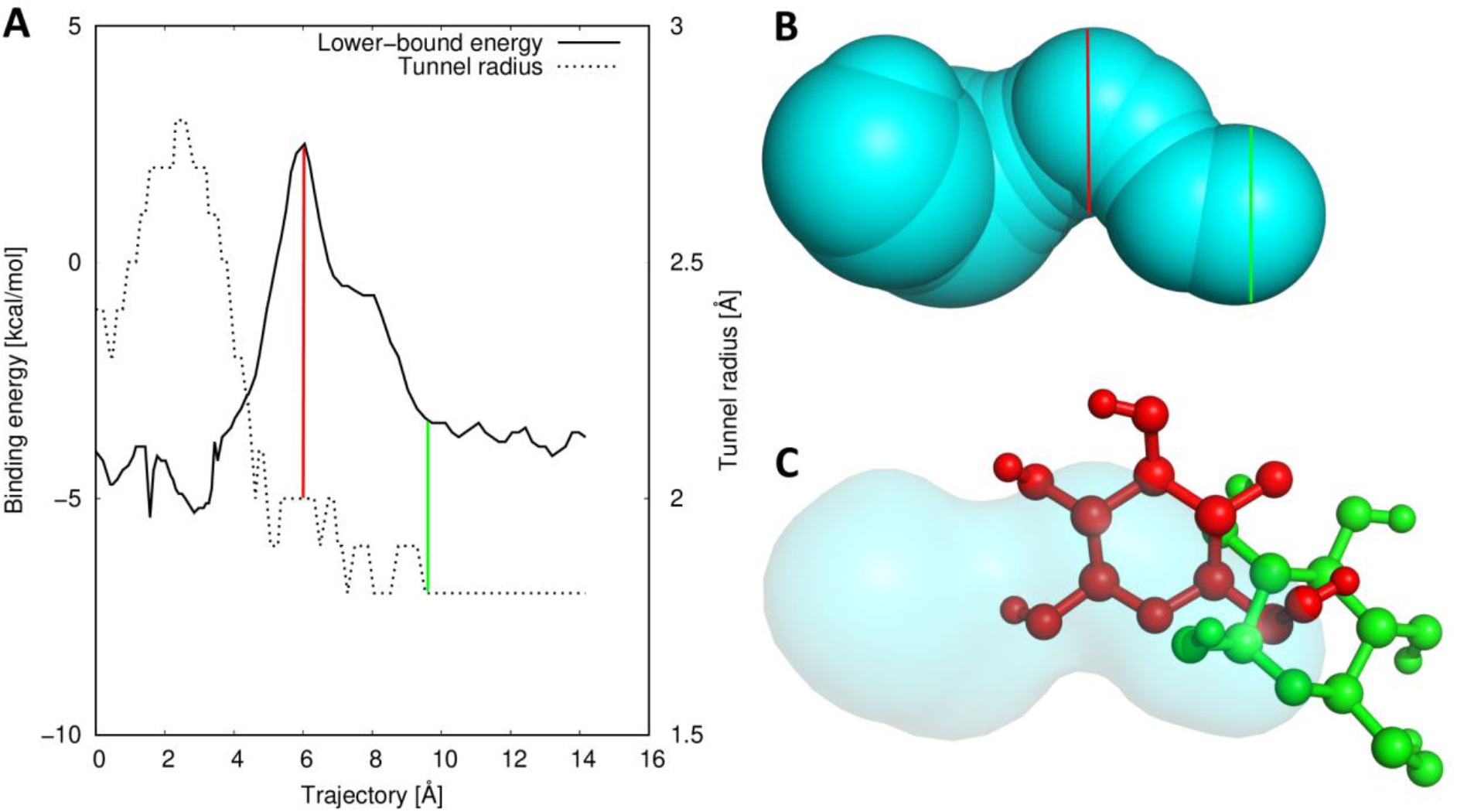
An example of the case with a large difference between the energetical maximum identified by CaverDock and the geometrical bottleneck identified by CAVER. (A) Energy profile from CaverDock (solid) and the geometric profile from CAVER (dotted). The tunnel region with the energy maximum is highlighted with the red line, and the region with a geometric bottleneck is highlighted with the green line. (B) Visualisation of the tunnel with highlights corresponding to the energy profile. (C) Visualisation of the cognate ligand ß-D-glucose conformations extracted from the trajectory from tunnel 1 of the structure of glucose dehydrogenase (PDB ID 2VWG). The binding pose based on the energy maximum (red) and geometric tunnel bottleneck (green).

CaverDock energy profiles were used to analyse the ligand preference of tunnels based on the energy barriers. We compared the maximum energies in up to the three tunnels and selected the best one. In the 13,158 proteins with successful CaverDock calculations, tunnel 1 had the most favourable energy in 75 % of the cases and tunnel 2 in another 19 % of the cases (Table S10). Therefore, for screening purposes, the analysis of tunnel 1 (or at most tunnel 2) would be enough for more than 93 % of proteins. Based on these results, tunnel 1 had the best properties for ligand un/binding and would be the most biochemically relevant.

Finally, we studied how well the cognate ligands were recognised by their receptors. We analysed the distribution of energy maxima (Figure S13) and the energy barrier (Figure S14). In the case of tunnel 1, which was the most preferred tunnel for ligand un/binding, almost 80 % of the E_Max_ values were in the range between −10 kcal/mol to 5 kcal/mol, and the energy barriers E_a_ were in the range between 0 kcal/mol to 10 kcal/mol. Both values were highly correlated for cognate ligands, and the Pearson’s correlation coefficient was 0.98 for energies from all three tunnels. Using E_Max_ seems to be equivalent to E_a_ for cognate ligands and probably for other natural substrates, which should be transported reasonably fast and bound in the active site. For inhibitors, both values could have different meanings as the molecule does not need to pass the entire way to the active site, but the binding affinity must be much stronger. In such cases, we recommend using E_Max_ values as they are easier to collect and interpret. We analysed the data split by pocket classes, EC numbers, and cognate ligand similarity. Still, all the datasets showed similar trends without major differences (data not shown), implying that ligand trajectories are case-specific rather than showing some general trends in different groups of enzymes.

## 4. Discussion and conclusions

We describe development of an automatic pipeline for the analysis of pockets and tunnels in enzymes and its application to study enzyme-cognate ligand complexes. The results provided a way to select potentially biologically relevant tunnels. The proposed approach can be used for extending large protein datasets for structural analyses and screenings. We analysed more than 17,000 cognate enzyme-ligand complexes. We were able to successfully annotate and analyse structural features and the energetics of ligand passage through tunnels in 13,158 enzyme structures. The data collected in this study will be made available to the wide scientific community via PDBe Knowledgebase ^52^. Each part of the pipeline was thoroughly validated, and the data showed that binding pockets selected based on the location of a bound ligand had a good overlap with catalytic and binding residue annotations from the structural databases. Therefore, bound ligands can be used to extend the datasets for pocket and tunnel analyses. Our experiments showed that selecting the pocket purely by score or druggability from FPOCKET would be significantly less precise. On the other hand, our pipeline is limited to enzyme structures with bound ligands, which limits its use. However this limitation is merely a consequence of being able to classify enzymes and their cognate ligands based on their reactions, which are all available in public databases. Extrapolation of ligand positions among homologous protein structures could remove this limitation for many structurally or functionally related proteins.

The presented machine learning predictor for the annotation of pockets has proven to be efficient in deciding on the type of pocket. The predictor can be used for selecting whether a particular enzyme and pocket are viable for tunnel calculations. The structural analyses revealed that it is possible to select potentially biologically relevant tunnels both in ligand-bound and ligand-free structures. Tunnels are present in the enzymes of all seven EC classes. Strikingly, the ligand transport calculations revealed that the energetic maximum was not in the geometrical bottleneck in 50 % of analysed tunnels. Therefore, energy profiling provides a highly relevant information about hot spots for enzyme engineering that could not be obtained by analysing the tunnel geometry. The comparison of CaverDock energetic maxima for calculated tunnels in each enzyme structure indicated that tunnel 1 had the lowest energy barrier in 75 % of cases. Therefore, the geometric analysis was not sufficient for identification of the most relevant tunnels in a quarter of all studied cases. To improve the predictive power of such analysis, the study of geometrical and energetical bottlenecks should be done on large set of dynamic snapshots. The results from single structure may be biased by the enzyme conformation in the crystal structure.

The knowledge and data acquired in this study will be important for future screening studies and the development of computational tools. We showed that the presented pipeline could be used to generate features for machine learning predictors and to provide valuable information for key repositories of biological data, like PDBe Knowledgebase ^52^. The validation of CaverDock against MD simulations proved that approximative methods are precise enough for fast energetical analyses of ligand passages. Approximative methods and enhanced sampling simulations are necessary to simulate ligand transport within reasonable times. Thus, we recommend energy calculations with approximative methods for protein engineering studies. Our comprehensive analysis of protein tunnels and the passages of cognate ligands let us formulate following recommendations, which are valuable for the protein engineering community:

1. For analysis of tunnels in enzymes, start with the literature search and exploration of databases to determine essential residues, identify the location of the binding pocket, and transport pathways, whenever possible.
2. The pocket(s) that contains the essential functional residues should be used preferably. In the systems with unknown essential residues, the pocket which contains a bound cognate ligand of the enzyme should be used. If there are no ligand-bound structures for the enzyme of interest, analyse available structures of homologous enzymes which contain the ligand. We recommend caution when selecting the binding pocket based solely on the predicted scores by the tools for pocket calculation.
3. The most important step of the tunnel analysis is to set the starting point correctly. When annotations of essential residues are not available the conserved residues are another possibility. Otherwise, we recommend using the residue inside of the selected pocket, closest to the centre of the biological unit or the analysed protein chain in the asymmetrical unit to start the tunnel calculation from the deep part of the pocket. An incorrectly set starting point may hinder the tunnel calculation and impact the geometry of found tunnels.
4. Selection of the biochemically relevant tunnel(s) should be preferably made based on the experimental literature data. When no such information is available, either focus on the first tunnel from a screening scenario or the first three tunnels according to the highest priority score. CAVER users are advised to inspect the tunnels with priority scores above 0.55. If none of the found tunnels has a priority score above this value, select a different starting point and redo the calculations.
5. If the starting point for tunnel calculation is selected correctly and the first tunnel is shorter than 5 Å, the binding pocket may well be located on the surface and tunnel analysis will might not be relevant.
6. Analysis of tunnels should be complemented by study of substrate or product passage whenever possible.
7. Use the ranges of energy barriers defined in this study to filter out molecules with poor (un)binding (E_Max_: −10 kcal/mol to 5 kcal/mol, E_a_: 0 kcal/mol to 10 kcal/mol) for energetic analyses of ligand passage by the approximative method CaverDock ^14^. Other methods available for this purpose are SLITHER ^53^, MoMA-LigPath ^54^, GPathFinder ^15^, and ART-RRT ^16^.
8. Binding and unbinding studies by the approximative methods can be significantly enhanced by the analysis of an ensemble of structures obtained even from a short molecular dynamics simulations.

## Supporting information

Details from analyses

Filtered input dataset

Information for 8 validation systems

PDB ID pairs of the complexes and ligand free structures

Predictor code with datasets and prediction results

## Acknowledgements

The authors thank Dr. Sérgio Marques for valuable advice during the design of the pipeline and set up of validation simulations.

## Supplementary information

List of features and hyperparameters for the predictor; predictor learning curves; setup of ASMD simulations; detailed results from validations; details from structural and energetical analyses; tunnel parameters and presence in EC classes (PDF); filtered input dataset (CSV); information for 8 validation systems (CSV); list of PDB ID pairs of the complexes and ligand free structures (CSV); training dataset (CSV); testing dataset (CSV); predictor Python code (PY); training and testing dataset with labels from predictors (CSV).

