## Supplementary material for "Large-scale Annotation of Biochemically Relevant Pockets and Tunnels in Cognate Enzyme-Ligand Complexes": Details from analyses

[1] Loschmidt Laboratories, Department of Experimental Biology and RECETOX, Faculty of Science, Masaryk University, Kamenice 5, 625 00 Brno, Czech Republic; [2] International Clinical Research Center, St. Anne's University Hospital Brno, Pekařská 53, 656 91 Brno, Czech Republic; [3] European Molecular Biology Laboratory, European Bioinformatics Institute (EMBL-EBI), Wellcome Trust GenomeCampus, CB10 1SD Cambridge, United Kingdom

* Corresponding authors: S. Mazurenko and D. Bednar, Loschmidt Laboratories, Department of Experimental Biology and RECETOX, Faculty of Science, Masaryk University, Kamenice 5/A13, 625 00 Brno, Czech Republic; J. Thornton, European Molecular Biology Laboratory, European Bioinformatics Institute (EMBL-EBI), Wellcome Trust Genome Campus, CB10 1SD Cambridge, United Kingdom

**Keywords:** bottleneck, cognate ligand, cavity, enzyme, tunnel, machine learning, pocket, transport

Table S1: List of features used for training pocket annotation predictor

| Feature Origin | Feature |
|---|---|
| FPOCKET | Fpocket Score |
| | Druggability Score |
| | Number of Alpha Spheres |
| | Total Surface Area |
| | Polar Surface Area |
| | Apolar Surface Area |
| | Volume |
| | Mean local hydrophobic density |
| | Mean alpha sphere radius |
| | Mean alpha sphere solvent access |
| | Proportion of apolar alpha spheres |
| | Hydrophobicity score |
| | Volume score |
| | Polarity score |
| | Charge score |
| | Proportion of polar atoms |
| | Density of the cavity |
| | Maximum distance between two alpha spheres |
| | Bfactor score |
| Custom | Exposed ratio |

Table S2: List of hyperparameters of algorithms and their values

| Algorithms | Hyperparameters | Values | | Tested ranges |
|---|---|---|---|---|
| | | Two-class dataset | Three-class dataset | |
| KNN | K (Number of neighbors) | 17 | 6 | 1..30 |
| Random Forest | max_depth (maximum depth of trees) | 2 | 2 | 1..5 |
| | n_estimators (number of trees) | 4 | 4 | 1..5 |
| SVM | Kernel | Linear | Linear | ['rbf', 'poly', 'linear'] |
| | C (regularization parameter) | 0.13 | 0.14 | -3..0 (Logarithm with 50 steps) |
| Shallow ANN | Alpha (L2 penalty) | 0.1 | 1 | -1..0 (Logarithm with 10 steps) |
| | Hidden_layer_sizes (number of hidden layer neurons) | 43 | 21 | 2..64 |
| | Activation function | tanh | tanh | ['identity', 'logistic', 'tanh', 'relu'] |
| | Solver | sgd | sgd | ['lbfgs', 'sgd', 'adam'] |

Table S3: The structure of the training and test data based on the labels.

| | # Data points | # Surface (1) | # Borderline (0) | # Buried (-1) |
|---|---|---|---|---|
| **Train set** | 200 | 92 | 58 | 50 |
| **Test set** | 100 | 40 | 30 | 30 |

Table S4: Validation systems and applied settings for simulations.

| PDB-ID | Tunnel | ASMD settings | | CaverDock settings |
|---|---|---|---|---|
| | | Protein residue | Ligand atom | Drag atom |
| System #1 (PDB ID 1OTW) | 1 | A MET 221 | C9 | 26 |
| | 2 | A LYS 214 | | |
| | 3 | A THR 73 | | |
| System #2 (PDB ID 2BFN) | 1 | GLY 18 | C2 | 1 |
| | 2 | ARG 191 | | |
| | 3 | LEU 177 | | |
| System #3 (PDB ID 2RFY) | 1 | ARG 129 | O1 | default |
| | 2 | PRO 249 | | |
| | 3 | THR 248 | | |
| System #4 (PDB ID 2UWH) | 1 | THR 269 | C12 | 10 |
| | 2 | ASP 69 | | |
| | 3 | LEU 182 | | |
| System #5 (PDB ID 4E2Z) | 1 | HIS 181 | N | default |
| | 2 | ILE 390 | | |
| | 3 | GLY 138 | | |
| System #6 {PDB ID 5EDT) | 1 | THR 243 | C7 | default |
| | 2 | ALA 349 | | |
| | 3 | HEME | | |
| | 4 | LEU 209 | | |
| System #7 (PDB ID 3ORW) | 1 | LEU 130 | C1 | 12 |
| | 2 | ALA 251 | | |
| System #8 (PDB ID 5U6M) | 1 | ASN 346 | P1 | default |
| | 2 | PHE 354 | C | |
| | 3 | ARG 28 | C6 | |

Table S5: Percentages of pockets with the best FPOCKET and druggability scores.

| Subset | Pocket type based on ligand coverage | Number of cases | Ligand-binding pocket has best FPOCKET score [%] | Ligand-binding pocket has best Druggability score [%] |
|---|---|---|---|---|
| **Pockets with matching annotations** | All pockets | 8,350 | 42.83 | 22.50 |
| | Single pocket | 3,344 | 49.85 | 25.48 |
| | One main pocket | 4,262 | 39.32 | 21.52 |
| | Two main pockets | 744 | 31.32 | 14.78 |
| **Pockets with no matching residue annotations** | All pockets | 2,696 | 26.97 | 12.20 |
| | Single pocket | 941 | 30.50 | 14.45 |
| | One main pocket | 1,464 | 26.98 | 10.93 |
| | Two main pockets | 291 | 15.46 | 11.34 |
| **Pockets with no annotations** | All pockets | 4,651 | 40.94 | 17.37 |
| | Single pocket | 1,682 | 50.65 | 20.15 |
| | One main pocket | 2,528 | 37.46 | 16.30 |
| | Two main pockets | 441 | 23.81 | 12.93 |

Table S6: The performance of five selected predictors on training and test data. The reported values on the training set are based on the 5-fold cross-validation. All the values are given as averages and standard deviations based on 100 independent runs.

| Dataset | Method | Accuracy | | F1 | | 1-FPR | |
|---|---|---|---|---|---|---|---|
| | | Training set | Test set* | Training set | Test set* | Training set | Test set* |
| Three-class problem | KNN (k=6) | 0.51 ± 0.05 | 0.47 | 0.47 ± 0.05 | 0.45 | 0.70 ± 0.12 | **0.75** |
| | SVM | **0.56 ± 0.05** | 0.52 | 0.49 ± 0.05 | 0.44 | 0.69 ± 0.11 | 0.65 |
| | Random Forest | 0.48 ± 0.03 | 0.48 ± 0.04 | 0.51 ± 0.04 | 0.40 ± 0.05 | 0.43 ± 0.13 | 0.37 ± 0.16 |
| | ANN (21 neurons) | 0.55 ± 0.02 | **0.54 ± 0.02** | **0.63 ± 0.02** | **0.50 ± 0.03** | 0.70 ± 0.04 | 0.67 ± 0.05 |
| | Naïve Bayesian | 0.34 ± 0.05 | 0.40 | 0.30 ± 0.05 | 0.19 | **0.90 ± 0.10** | 0.0 |
| Two-class problem | KNN (k=17) | 0.67 ± 0.04 | **0.71** | 0.65 ± 0.04 | **0.71** | 0.79± 0.05 | 0.70 |
| | SVM | **0.70 ± 0.04** | 0.70 | 0.69 ± 0.04 | 0.70 | 0.70 ± 0.06 | 0.62 |
| | Random Forest | 0.61 ± 0.03 | 0.66 ± 0.04 | 0.72 ± 0.02 | 0.65 ± 0.04 | 0.78 ± 0.06 | 0.68 ± 0.07 |
| | ANN (43 neurons) | 0.66 ± 0.02 | 0.70 ± 0.02 | **0.74 ± 0.02** | 0.70 ± 0.02 | 0.76 ± 0.02 | **0.70 ± 0.03** |
| | Naïve Bayesian | 0.57 ± 0.04 | 0.41 | 0.45 ± 0.05 | 0.41 | **0.93 ± 0.05** | 0.02 |

*KNN, SVM and Naïve Bayesian did not have any randomization, which we could use to have the standard deviation on the test set.

Table S7: CaverDock energies in static structures and snapshots from molecular dynamics.

| CaverDock energies in static structure | | | | Priority in ASMD | Averaged CD energies from 100 snapshots | | | |
|---|---|---|---|---|---|---|---|---|
| Case | Tunnel | Priority in PDB | $E_{max}$ [kcal/mol] | | Priority in MD | $E_{max}$ [kcal/mol] | Finished calculations | Tunnel occurrence* |
| System 1 (1OTW) | 1 | 1 | 20.4 | 2 | 3 | -1.3 | 13 | 13 |
| | 2 | 2 | 21.8 | 1 | 1 | 1.3 | 25 | 25 |
| | 3 | 3 | 46.6 | 3 | 5 | 15.4 | 8 | 8 |
| System 2 (2BFN) | 1 | 1 | -2.5 | 1 | 1 | -1.9 | 95 | 100 |
| | 2 | 3 | 1.7 | 2 | 2 | -1.2 | 78 | 78 |
| | 3 | 4 | 6.2 | 3 | None | None | 0 | 0 |
| System 3 (2RFY) | 1 | 1 | -3.3 | 1 | 1 | -3.5 | 61 | 69 |
| | 2 | 2 | -5.5 | 3 | 3 | -0.9 | 56 | 60 |
| | 3 | 3 | -1.9 | 2 | 2 | -4.6 | 58 | 62 |
| System 4 (2UWH) | 1 | 1 | -4.9 | 1 | 3 | -3.5 | 81 | 85 |
| | 2 | 2 | 2.8 | 2 | 1 | -4.1 | 78 | 79 |
| | 3 | 4 | 3.9 | 3 | 6 | -0.2 | 16 | 16 |
| System 5 (4E2Z) | 1 | 1 | -1.4 | 1 | 1 | -4.7 | 73 | 74 |
| | 2 | 2 | 7.8 | 2 | 2 | 1.2 | 84 | 89 |
| | 3 | 5 | 48.0 | 3 | 30 | 15.4 | 4 | 4 |
| System 6 {5EDT) | 1 | 1 | 28.7 | 2 | 1 | 10.5 | 60 | 60 |
| | 2 | 2 | 47.0 | 1 | 4 | 24.3 | 26 | 26 |
| | 3 | 3 | 19.2 | 3 | 3 | 15.1 | 47 | 47 |
| | 4 | 5 | 39.6 | 4 | 8 | 22.5 | 15 | 15 |
| System 7 (3ORW) | 1 | 1 | -0.6 | 1 | 1 | -3.8 | 96 | 96 |
| | 2 | 3 | 12.0 | 2 | 5 | 10.5 | 9 | 9 |
| System 8 (5U6M) | 1 | 1 | -6.2 | 1 | 1 | -5.4 | 90 | 92 |
| | 2 | 2 | -2.8 | 2 | 4 | -4.2 | 87 | 89 |
| | 3 | 4 | 31.1 | 3 | 5 | 12.4 | 33 | 33 |

* number of snapshots out of 100 containing the tunnel

Table S8: Length of the first three tunnels in predicted classes.

| Tunnel number | Predicted pocket class | Cases | Tunnel length | | |
|---|---|---|---|---|---|
| | | | x < 5 Å [%] | 5 Å < x < 15 Å [%] | 15 Å < x [%] |
| Tunnel 1 | Buried | 3,552 | 25.11 | 45.97 | 28.91 |
| | Borderline | 3,178 | 41.19 | 49.06 | 9.75 |
| | Surface | 7,702 | 74.24 | 22.44 | 3.32 |
| Tunnel 2 | Buried | 3,215 | 2.33 | 40.34 | 57.33 |
| | Borderline | 2,798 | 2.64 | 61.62 | 35.74 |
| | Surface | 6,142 | 6.61 | 70.55 | 22.84 |
| Tunnel 3 | Buried | 2,860 | 0.59 | 20.80 | 78.60 |
| | Borderline | 2,268 | 0.35 | 35.89 | 63.76 |
| | Surface | 4,907 | 1.35 | 44.37 | 54.29 |

Table S9: Match between energetic maximum and geometrical bottleneck.

| Tunnels | Cases | $E_{Max}$ in tunnel bottleneck | | | |
|---|---|---|---|---|---|
| | | Bottleneck disc [%] | Bottleneck area 1.5 Å [%] | Bottleneck area 3 Å [%] | Bottleneck area 5 Å [%] |
| Tunnels 1-3 | 29 693 | 50.35 | 63.91 | 70.39 | 74.49 |
| Tunnel 1 | 12 774 | 69.65 | 77.15 | 80.37 | 82.25 |
| Tunnel 2 | 9 449 | 39.68 | 55.54 | 63.51 | 68.41 |
| Tunnel 3 | 7 470 | 30.86 | 51.85 | 62.05 | 68.90 |

*data was collected from energy profiles with 33 % deleted parts

Table S10: Comparison of first three tunnels based on the energetic maximum.

| Type of data | Number of cases |
|---|---|
| Protein-ligand pairs for CaverDock calculations | 14,432 |
| Successful CaverDock calculations | 13,158 |
| Failed CaverDock calculations | 1,274 |
| Tunnel 1 is the best tunnel in comparison [%] | **74.75** |
| Tunnel 2 is the best tunnel in comparison [%] | 18.52 |
| Tunnel 3 is the best tunnel in comparison [%] | 6.73 |

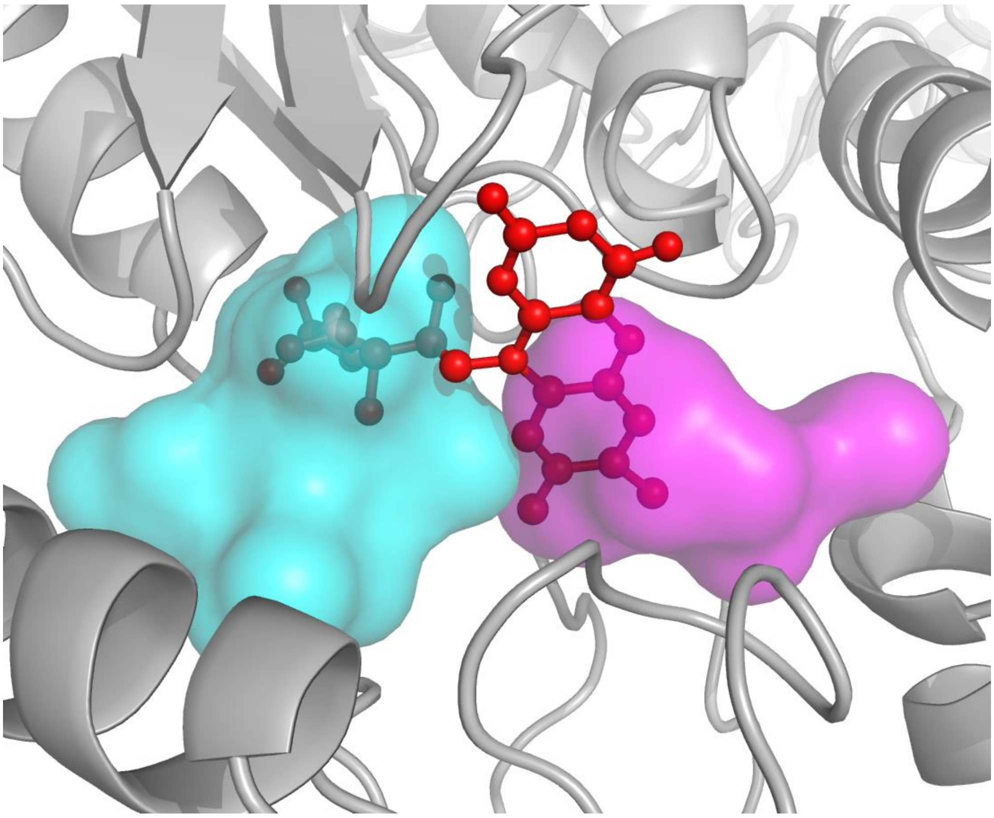

Figure S1: Visualisation of a flavin mononucleotide molecule, which is part of two main pockets in the structure of oxidoreductase FprA (PDB ID 1YCH). The first pocket (cyan) contains 55 % of the atoms of the bound ligand (red balls and sticks), and the second pocket (magenta) contains 48 % of the atoms. We illustrate how the settings for FPOCKET caused the algorithm to define the empty space as two pockets. Due to the low difference in ligand coverage, we selected pocket 1 as the main one based on the druggability score. This pocket does not contain any annotated residues, therefore, we did not have any correct matches.

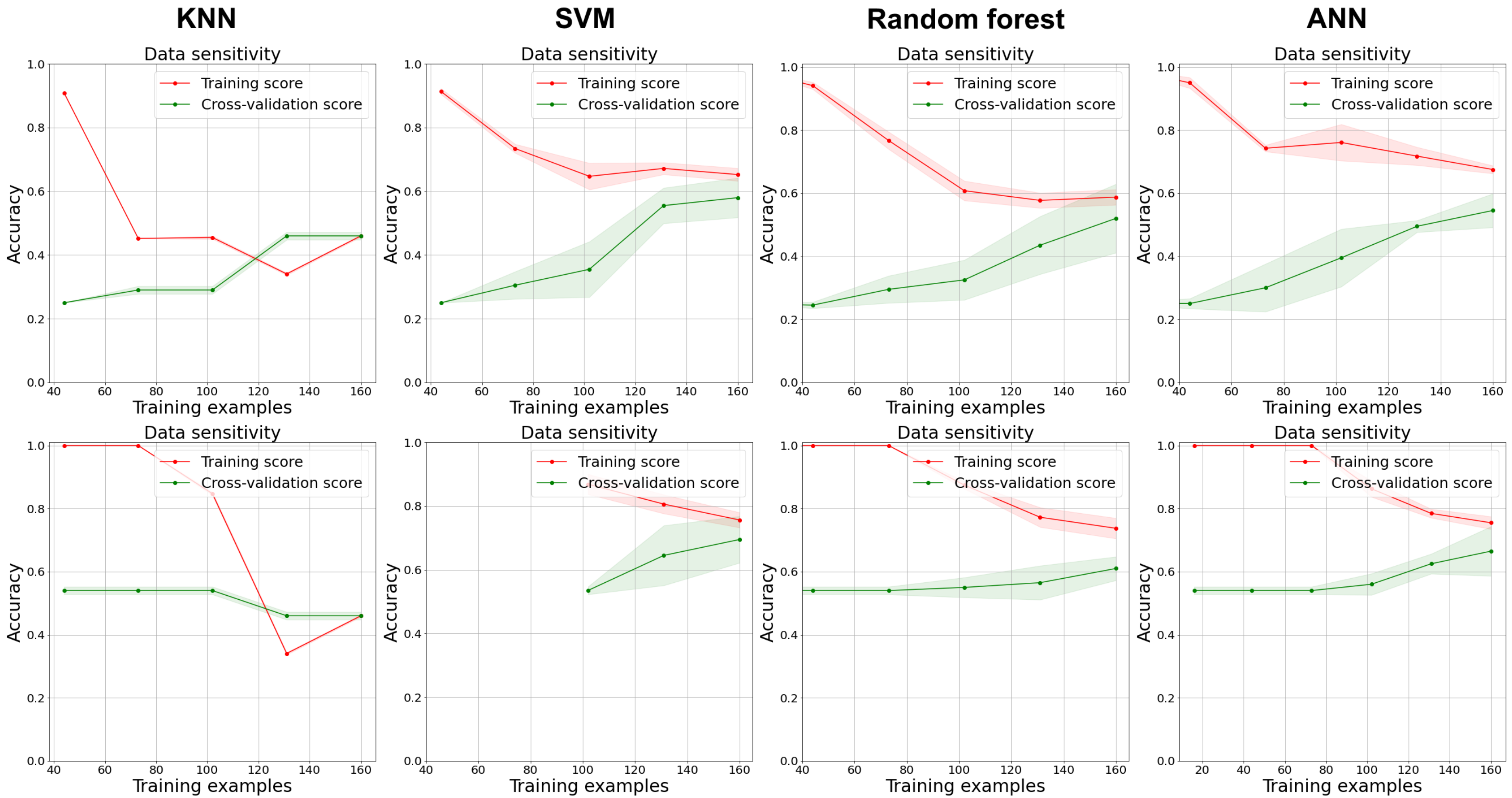


Figure S2: Learning curves for the predictors for the three-class (top) and two-class (bottom) problems. For KNN, NN and SVMs, the two curves have converged, indicating a good fit but also that adding more data is unlikely to improve the accuracy. For the Random Forest, the gap between training and cross-validation curves shows slight overfitting but indicates a trend towards similar values of 50% and 70% for the accuracy for three- and two-class problems, respectively.

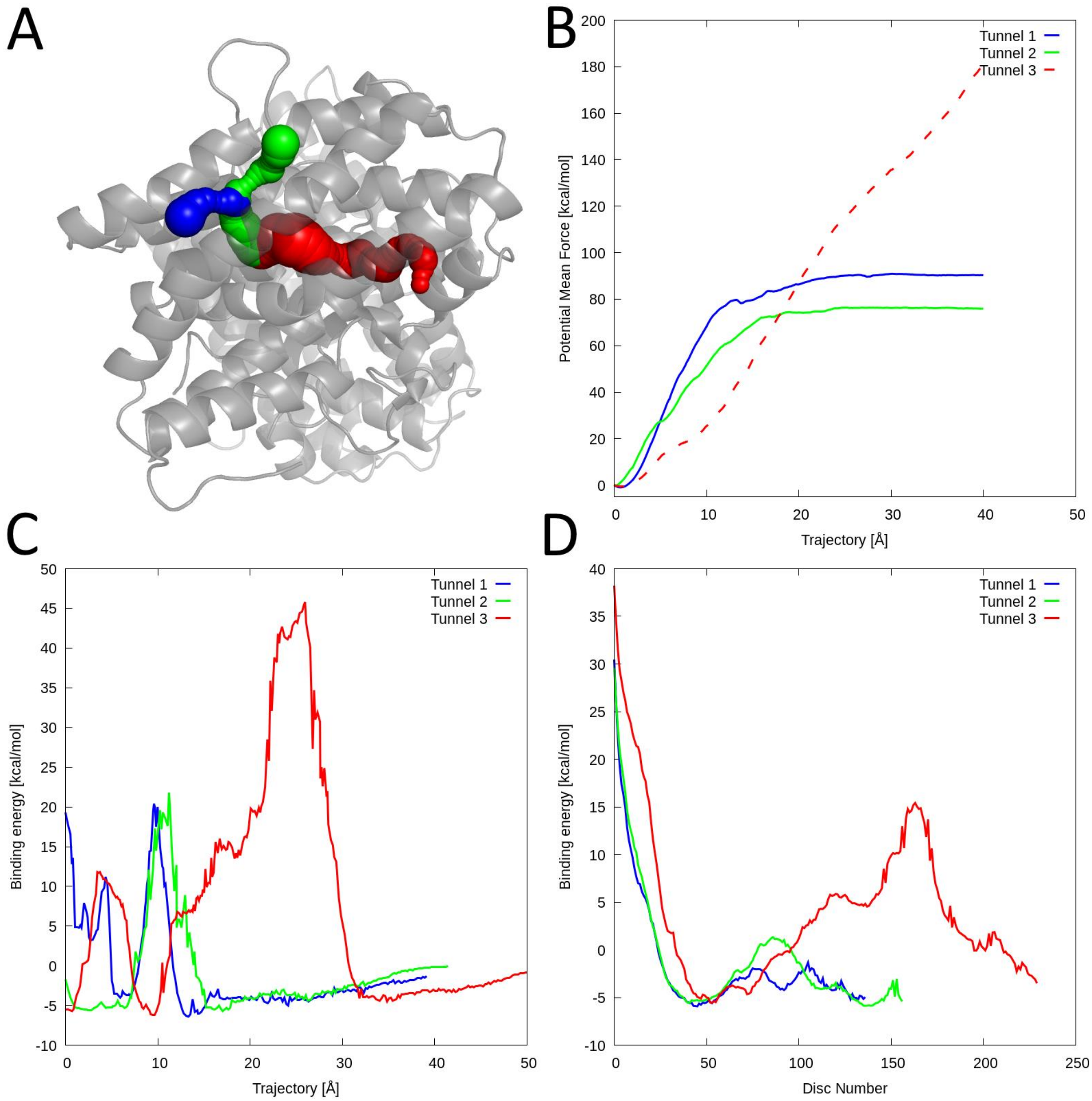


Figure S3: Results from CaverDock validation for System #1 - pyrroloquinoline-quinone synthase with pyrrolo-quinoline quinone. (A) Visualisation of the protein structure (PDB ID 1OTW) with analysed tunnels showed as spheres: tunnel 1 (blue), tunnel 2 (green), tunnel 3 (red); (B) Potential mean force (PMF) profiles from ASMD simulations, the profile for stuck ligand is shown as dashed line; (C) energy profiles from static CaverDock calculations; (D) averaged CaverDock energy profiles from 50 ns simulation snapshots.

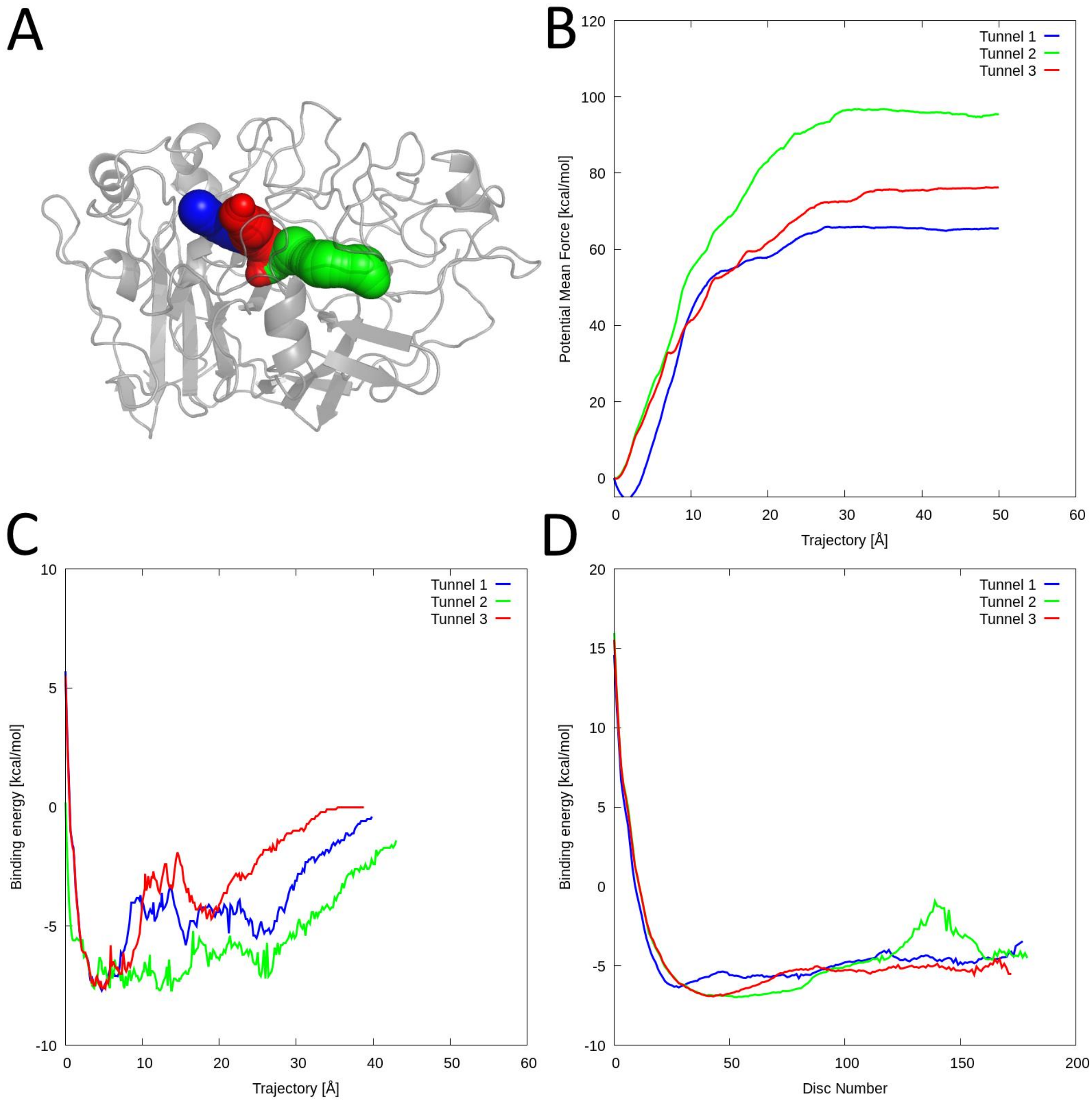


Figure S4: Results from CaverDock validation for System #3 - cellobiohydrolase with cellobiose. (A) Visualisation of the protein structure (PDB ID 2RFY) with analysed tunnels showed as spheres: tunnel 1 (blue), tunnel 2 (green), tunnel 3 (red); (B) PMF profiles from ASMD simulations; (C) energy profiles from static CaverDock calculations; (D) averaged CaverDock energy profiles from 50 ns simulation snapshots.

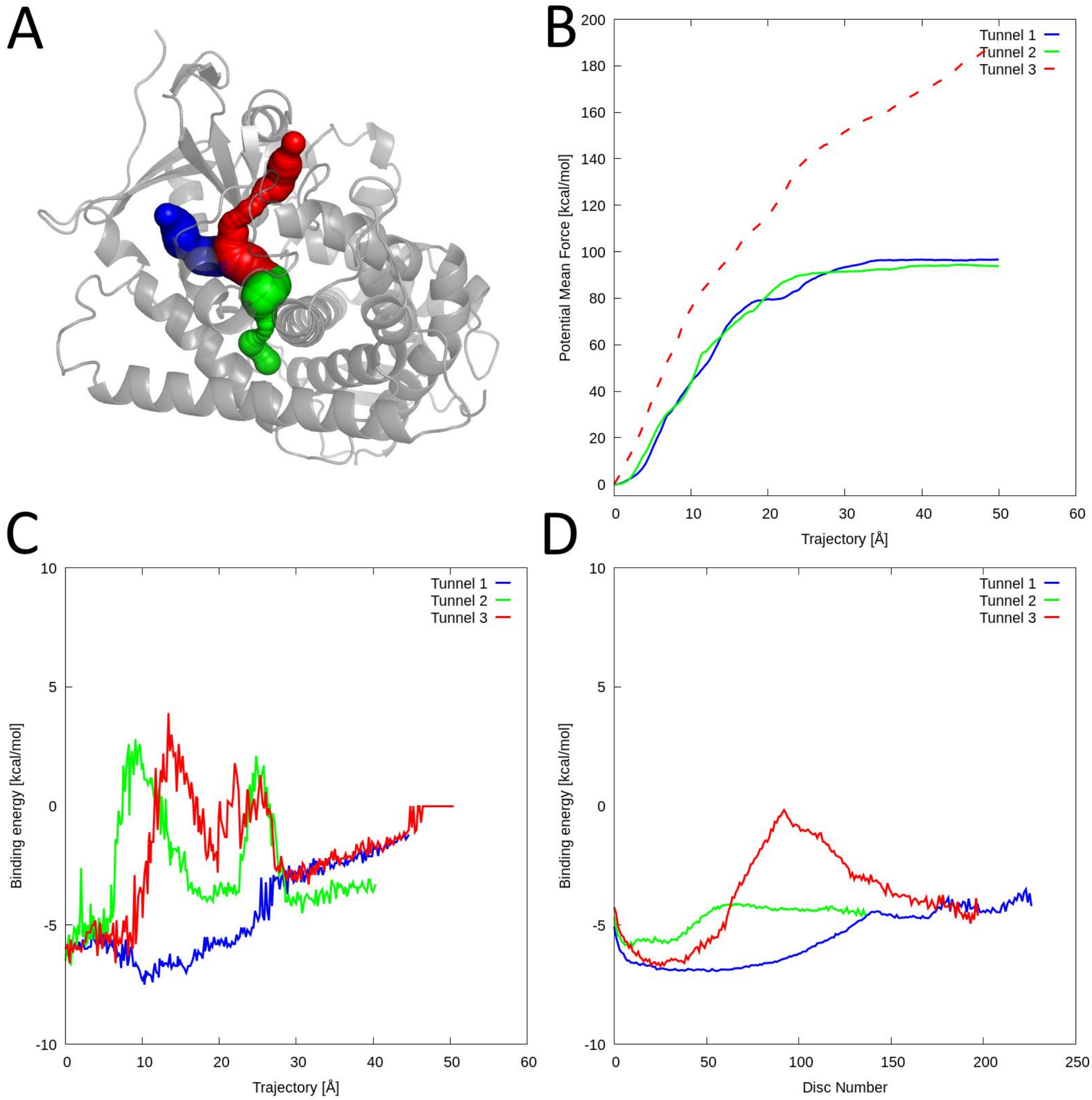


Figure S5: Results from CaverDock validation for System #4 - Cytochrome P450 BM3 with 11,14,15-trihydroxyicosatrienoic acid. (A) Visualisation of the protein structure (PDB ID 2UWH) with analysed tunnels showed as spheres: tunnel 1 (blue), tunnel 2 (green), tunnel 3 (red); (B) PMF profiles from ASMD simulations, the profile for stuck ligand is shown as dashed line; (C) energy profiles from static CaverDock calculations; (D) averaged CaverDock energy profiles from 50 ns simulation snapshots.

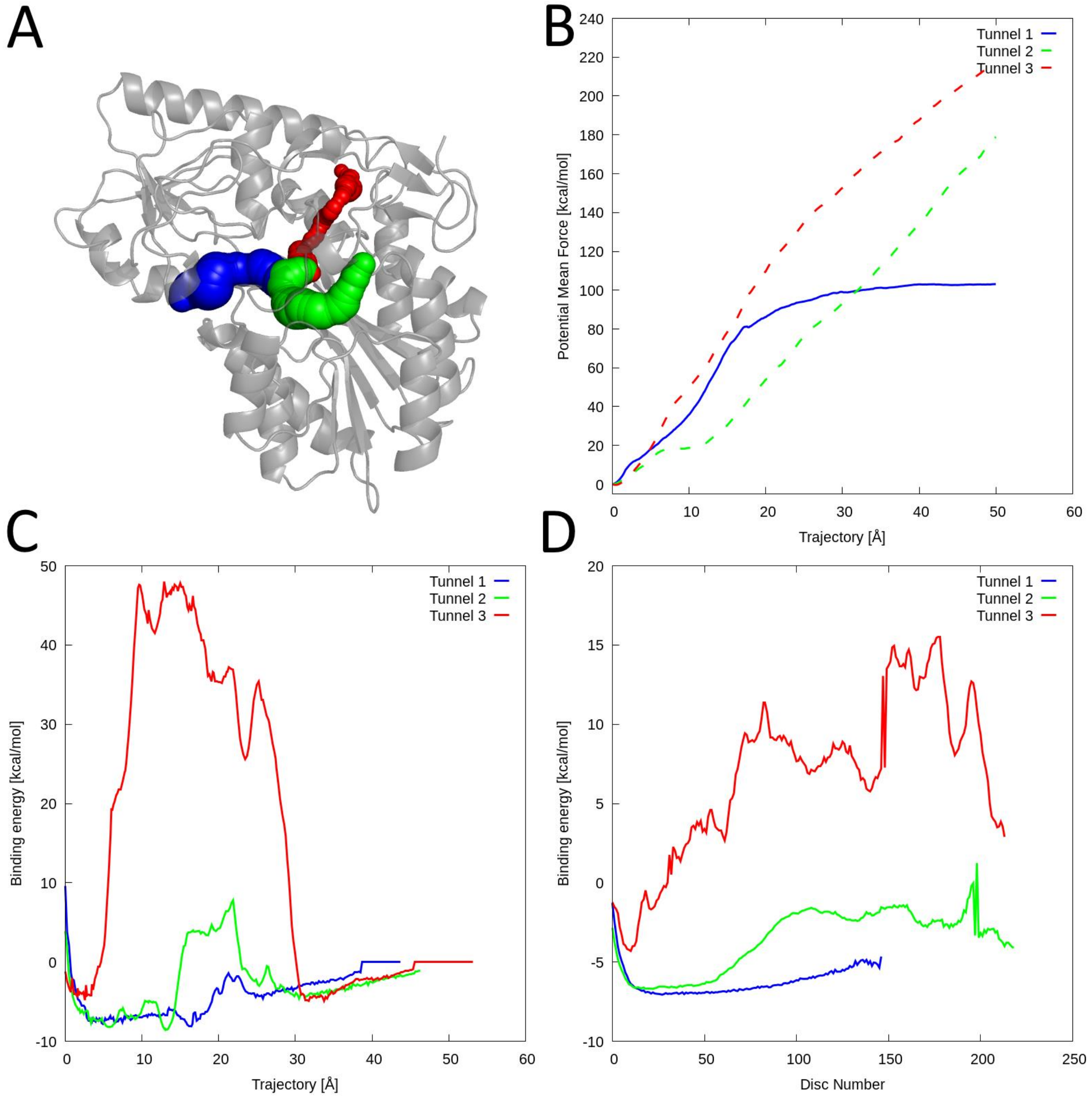


Figure S6: Results from CaverDock validation for System #5 - C-3'-methyltransferase with Se-adenosyl-L-selenohomocysteine. (A) Visualisation of the protein structure (PDB ID 4E2Z) with analysed tunnels showed as spheres: tunnel 1 (blue), tunnel 2 (green), tunnel 3 (red); (B) PMF profiles from ASMD simulations, the profiles for stuck ligand are shown as dashed line; (C) energy profiles from static CaverDock calculations; (D) averaged CaverDock energy profiles from 50 ns simulation snapshots.

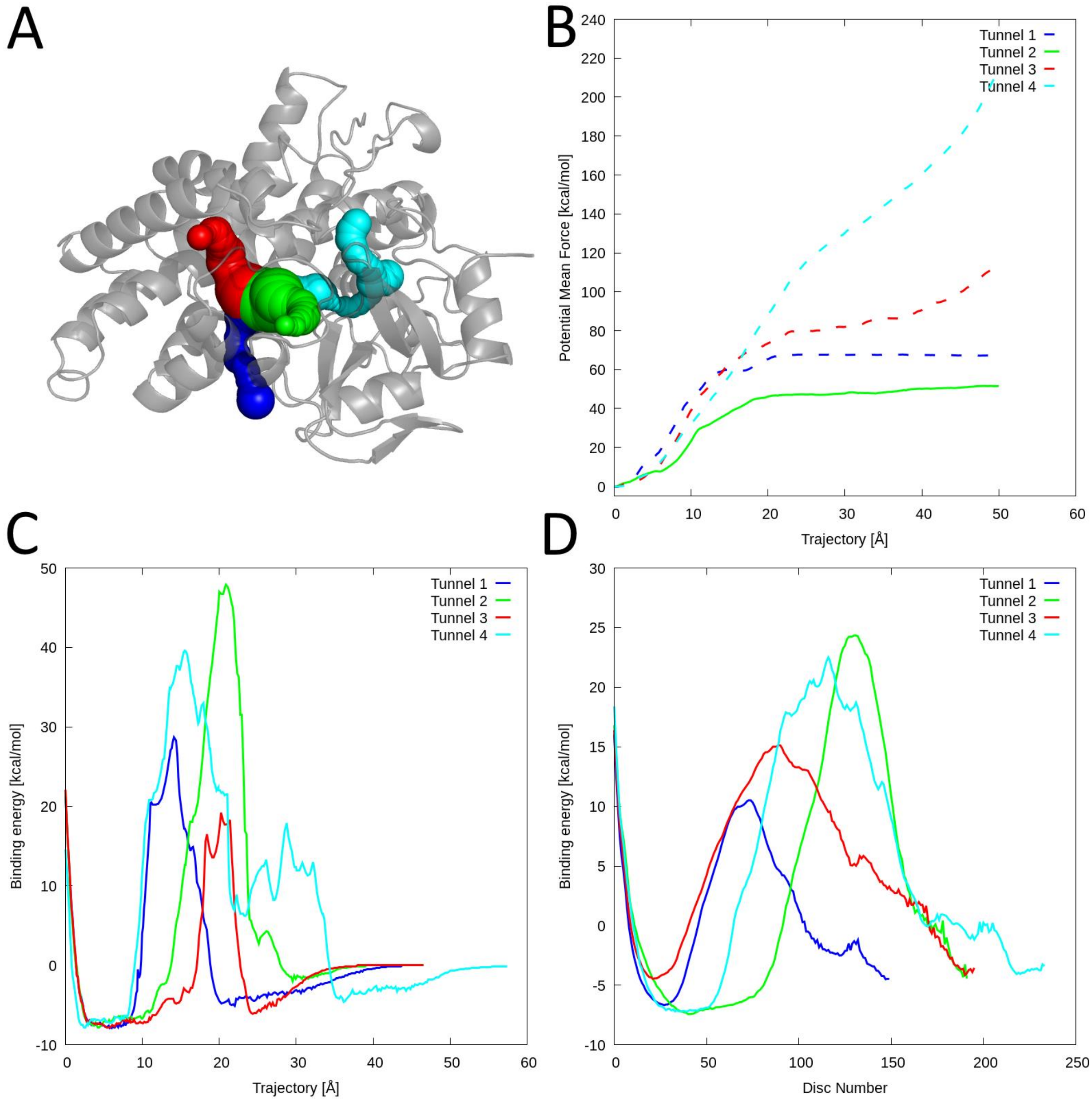


Figure S7: Results from CaverDock validation for System #6 - Cytochrome P450 CYP121 with (4*S*)-4-(5,5-Dimethylcyclohex-1-en-1-yl)cyclohex-1-ene-1-carboxylate. (A) Visualisation of the protein structure (PDB ID 5EDT) with analysed tunnels showed as spheres: tunnel 1 (blue), tunnel 2 (green), tunnel 3 (red), tunnel 4 (cyan); (B) PMF profiles from ASMD simulations; (C) energy profiles from static CaverDock calculations, the profiles where ligand unbound through incorrect tunnel are shown as dashed line; (D) averaged CaverDock energy profiles from 50 ns simulation snapshots.

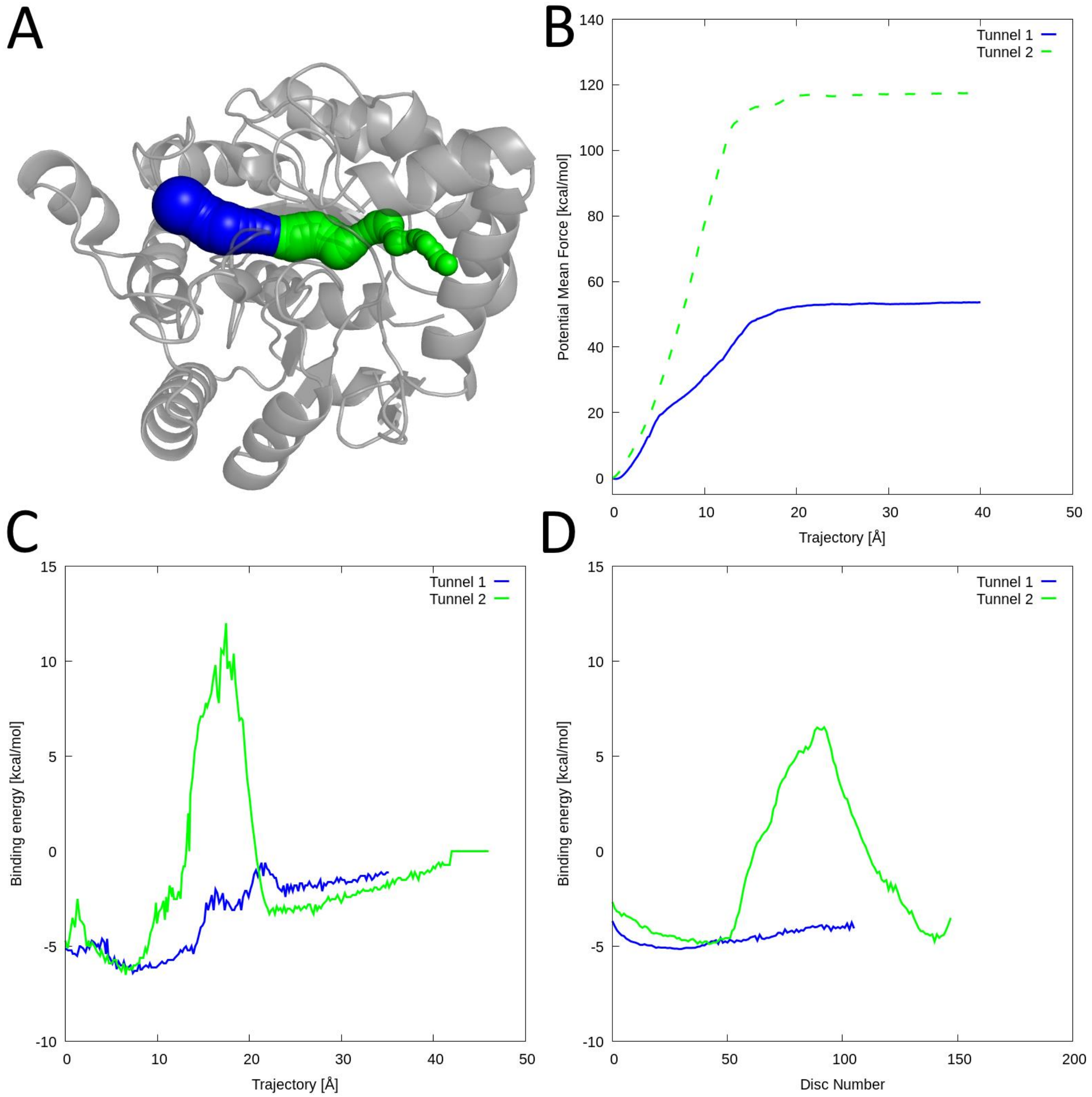


Figure S8: Results from CaverDock validation for System #7 – phosphotriesterase with N-(6-aminohexanoyl)-6-aminohexanoate. (A) Visualisation of the protein structure (PDB ID 3ORW) with analysed tunnels showed as spheres: tunnel 1 (blue), tunnel 2 (green); (B) PMF profiles from ASMD simulations, the profile where ligand unbound through incorrect tunnel is shown as dashed line; (C) energy profiles from static CaverDock calculations; (D) averaged CaverDock energy profiles from 50 ns simulation snapshots.

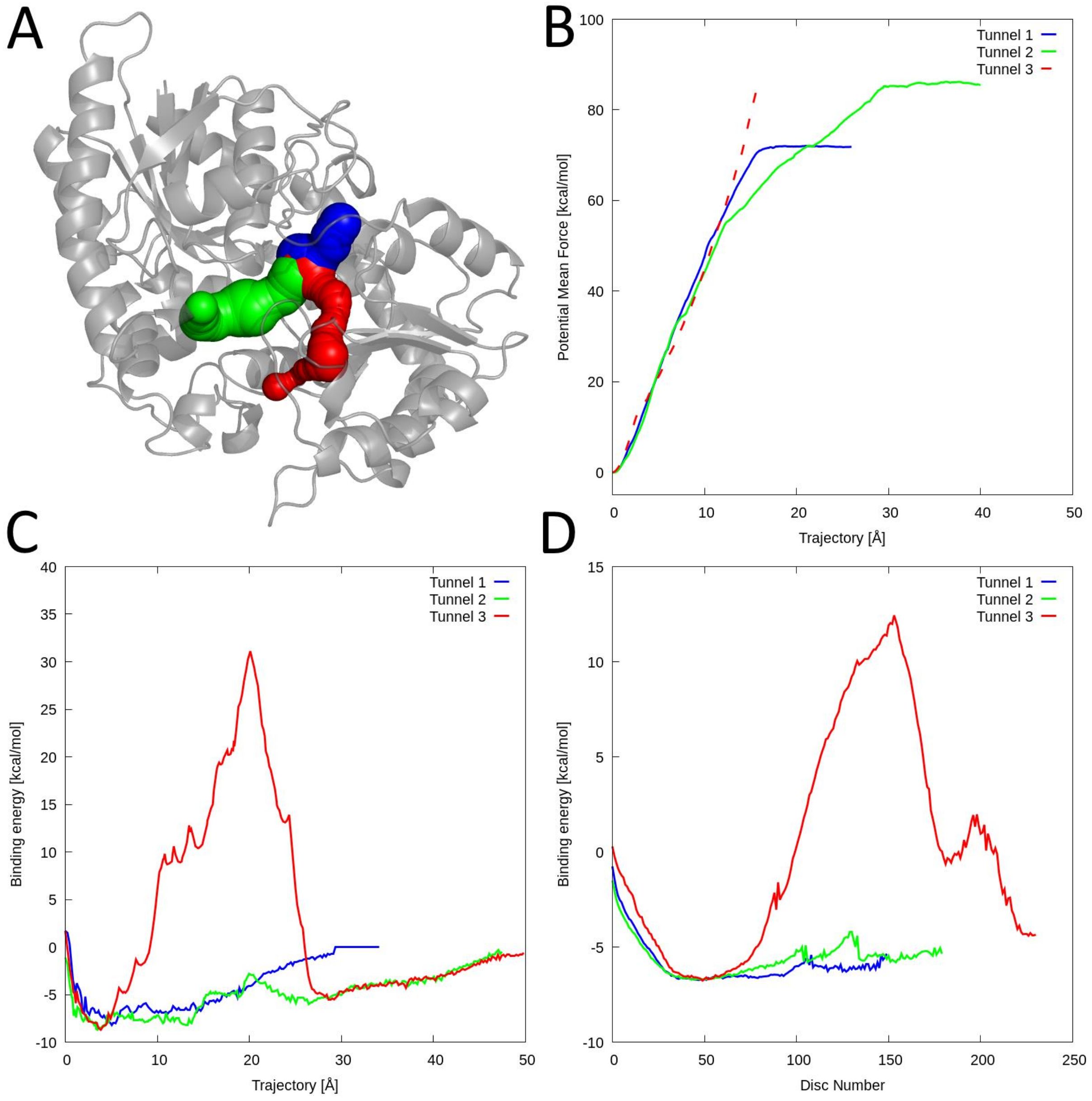


Figure S9: Results from CaverDock validation for System #8 – UDP-glucosyltransferase with uridine 5'-diphosphate. (A) Visualisation of the protein structure (PDB ID 5U6M) with analysed tunnels showed as spheres: tunnel 1 (blue), tunnel 2 (green); (B) PMF profiles from ASMD simulations, the profile for stuck ligand is shown as dashed line; (C) energy profiles from static CaverDock calculations; (D) averaged CaverDock energy profiles from 50 ns simulation snapshots.

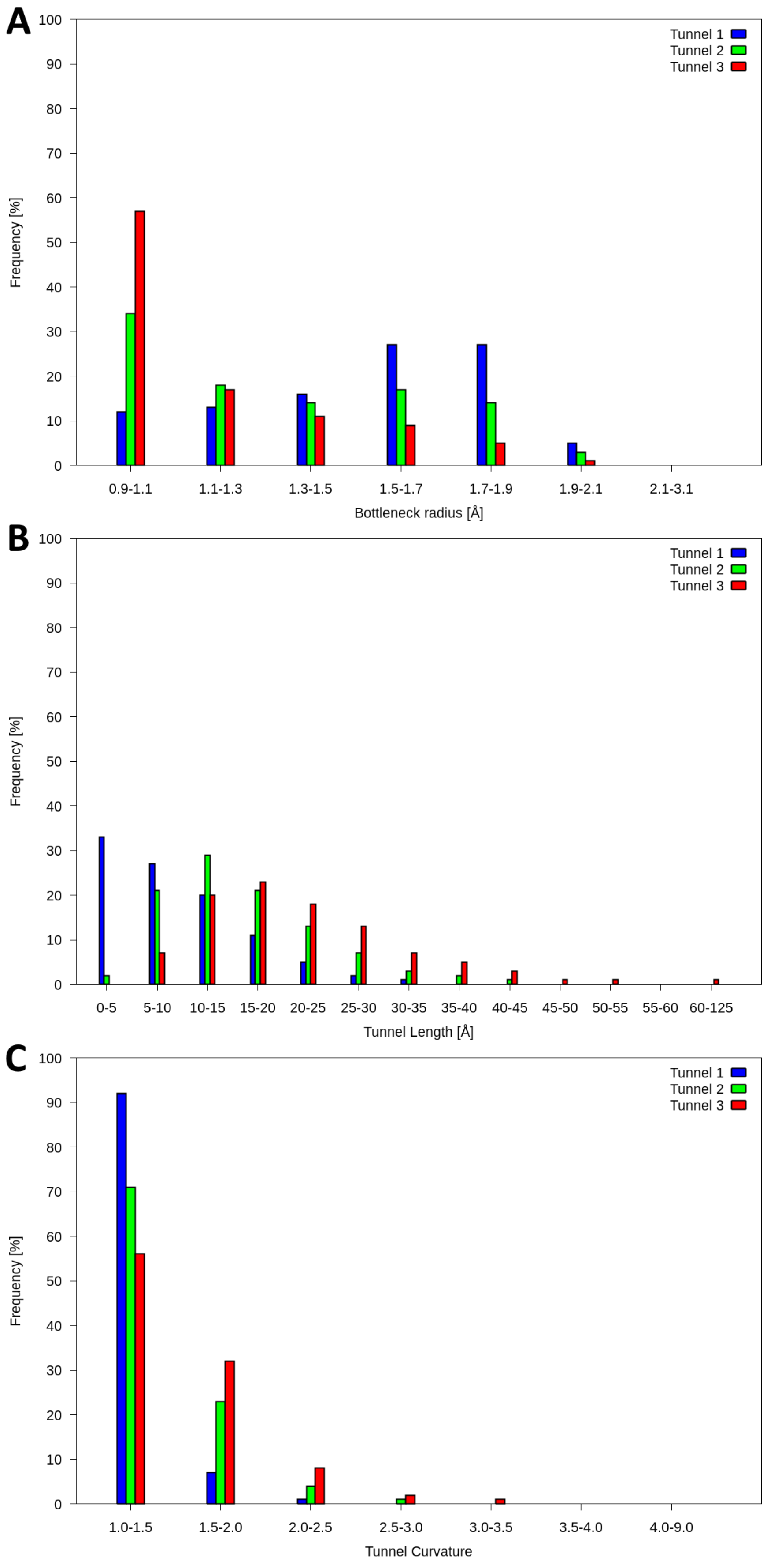


Figure S10: Distribution of tunnel parameters for first three tunnels from all tunnel cases (A) bottleneck radius, (B) tunnel length, (C) curvature.

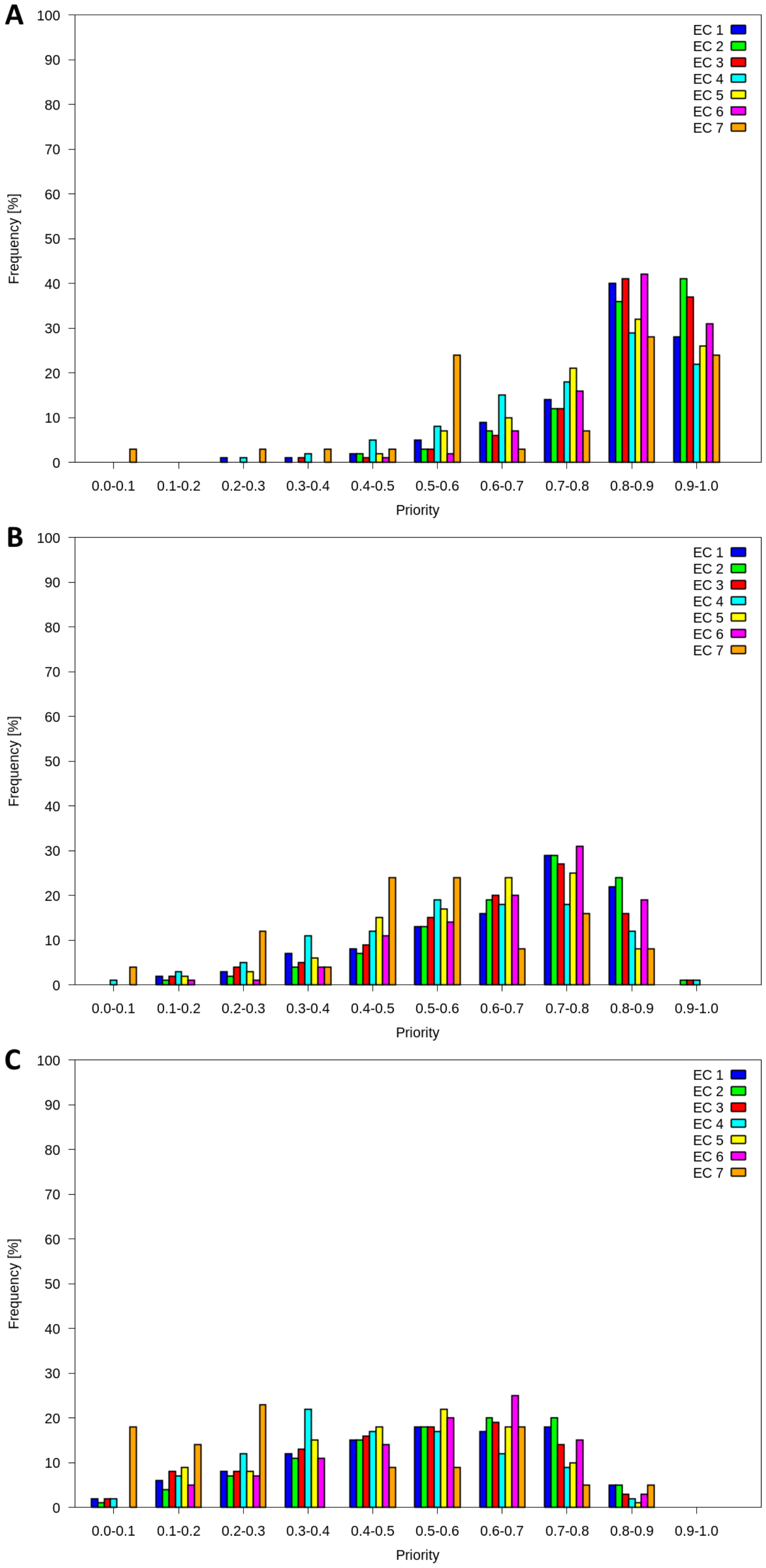


Figure S11: Distribution of tunnel priority scores in EC classes in the whole tunnel dataset for (A) tunnel 1, (B) tunnel 2, (C) tunnel 3.

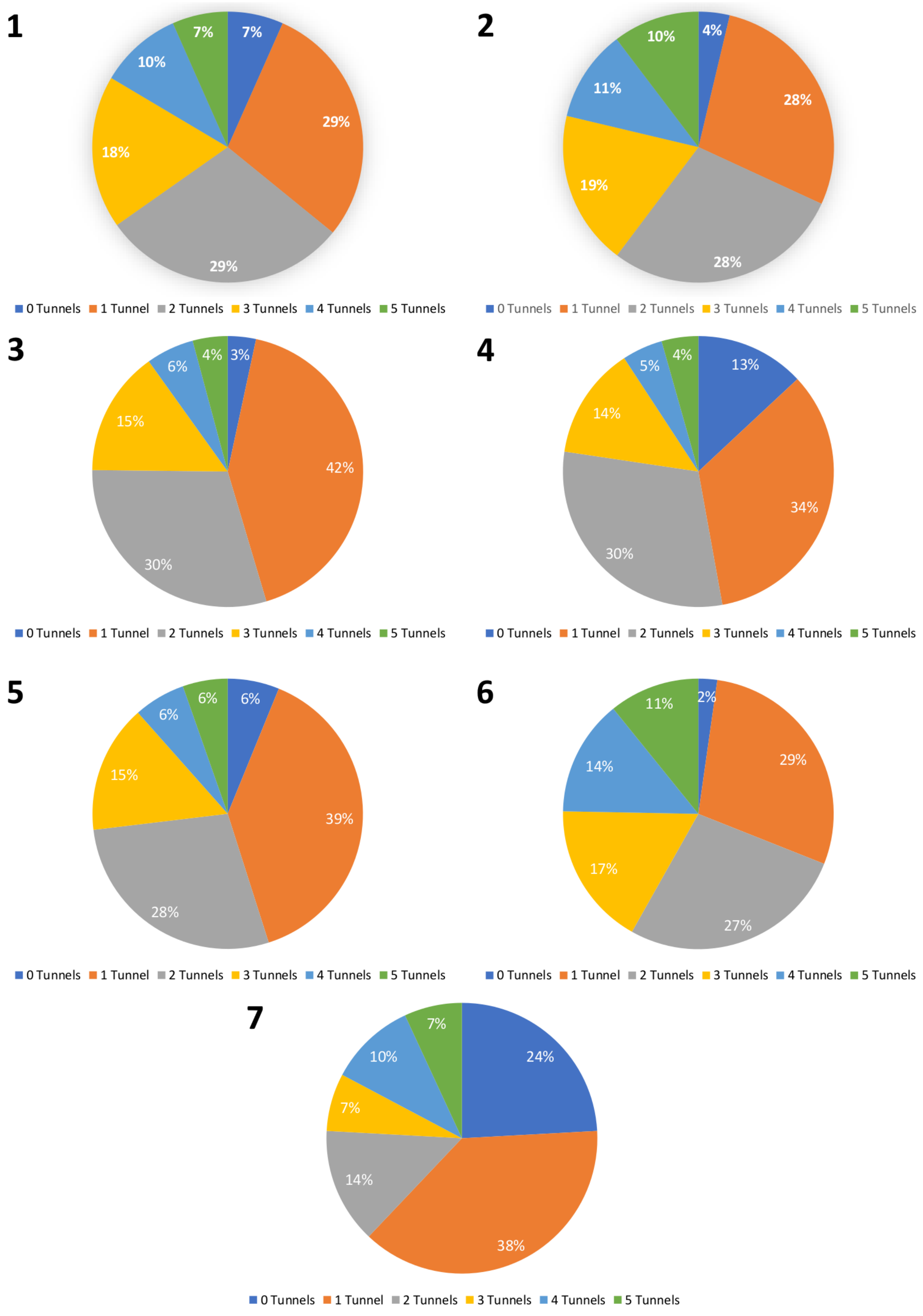


Figure S12: Number of tunnels with priority score above 0.55 found in tunnel dataset separated for each EC class (1-7).

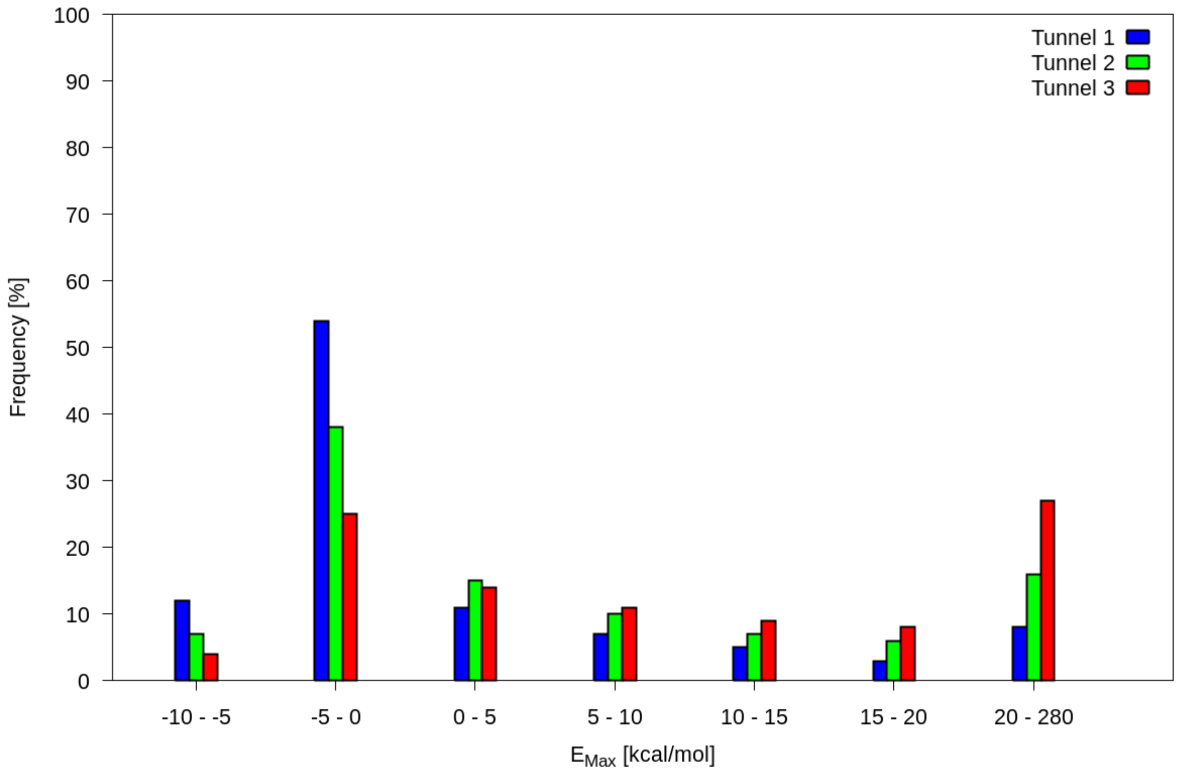


Figure S13: Distribution of energy maximum values in the CaverDock dataset for the first three tunnels. Three outliers with $E_{Max}$ energy around -50 kcal/mol were removed from the figure.

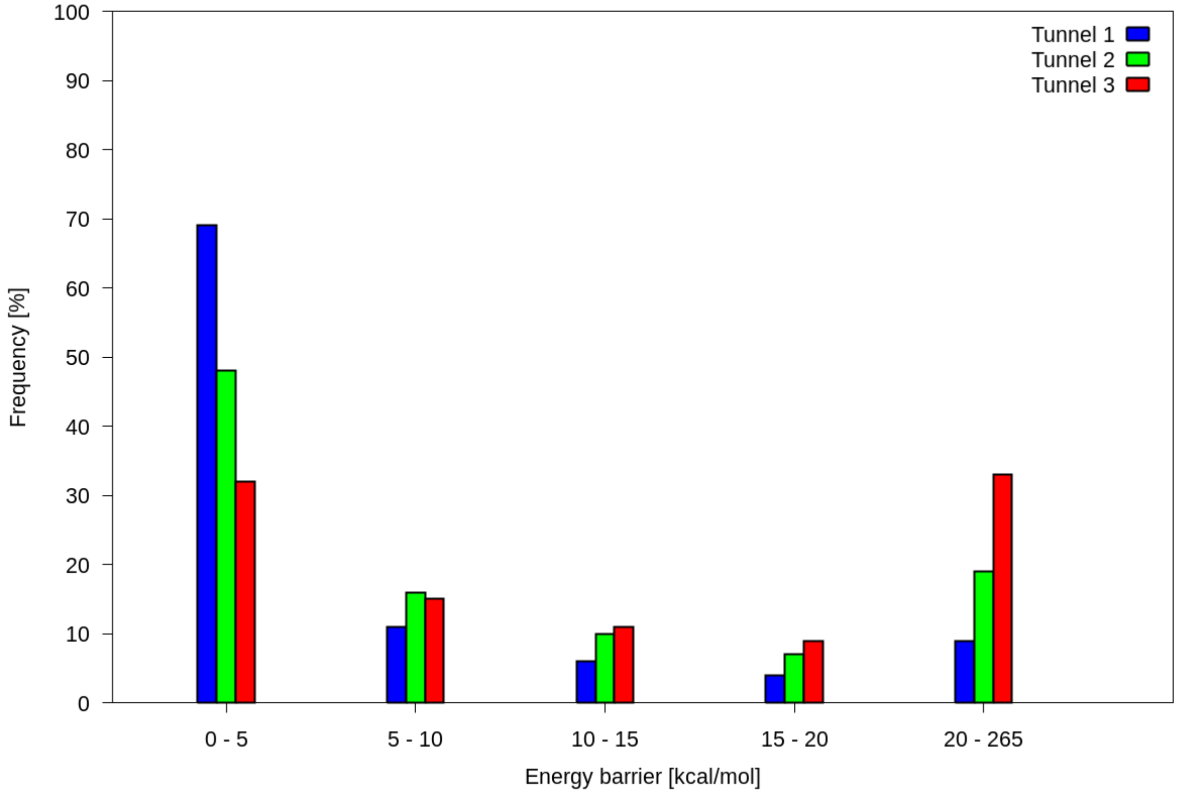


Figure S14: Distribution of energy barriers ($E_a$) in the CaverDock dataset for first three tunnels.
