## Supplementary material for "Large-scale Annotation of Biochemically Relevant Pockets and Tunnels in Cognate Enzyme-Ligand Complexes": Predictor code with datasets and prediction results: Final Code.py

```
# -*- coding: utf-8 -*-
"""TestLL.ipynb

Automatically generated by Colaboratory.

Original file is located at
    https://colab.research.google.com/drive/1YuGPR8u9nRh2x_WqRIcmgVaqzYPeea73

# Reading Data
"""

from google.colab import files
uploaded = files.upload()

from google.colab import files
uploaded_test = files.upload()

"""# Import some libraries"""

import pandas as pd
import numpy as np
from __future__ import print_function    
from keras.models import Sequential
import keras
from keras.models import Sequential
from keras.layers import Dense, Flatten, Conv1D
from keras.callbacks import ModelCheckpoint
from keras.models import model_from_json
from keras import backend as K
from sklearn.preprocessing import StandardScaler
import matplotlib.pyplot as plt
import seaborn as sns

"""# Dataframes"""

import io
df = pd.read_csv(io.BytesIO(uploaded['ML_subset2_with_seq.csv']))
print(df.head)
#df_test= pd.read_csv(io.BytesIO(uploaded_test['ML_subset-TESTING.csv']))

import io
df_test = pd.read_csv(io.BytesIO(uploaded_test['ML_subset-TESTING.csv']))
print(df_test)

"""# X - Y"""

X= df.iloc[ :, 1: 22]
#print(scaled_features)
Y= df ['Pocket Type']
print(Y)

X_test= df_test.iloc[ :, 1: 22]
print(X_test)
#print(scaled_features)
Y_test= df_test ['Pocket Type']
print(Y_test)

"""# Drop Ligand coverage"""

X_test=X_test.drop(columns=['Ligand coverage'])

X=X.drop(columns=['Ligand coverage'])

"""# Scaler"""

scaled_features = StandardScaler().fit_transform(X)

X_test = StandardScaler().fit_transform(X_test)

"""# KNN"""

from sklearn.model_selection import GridSearchCV
from sklearn.metrics import make_scorer, accuracy_score, precision_score, recall_score, f1_score
from sklearn.neighbors import KNeighborsClassifier
from sklearn.model_selection import cross_val_score
from sklearn.metrics import make_scorer, accuracy_score, precision_score, recall_score, f1_score
from sklearn.model_selection import KFold
from sklearn.model_selection import cross_validate
from sklearn.metrics import plot_confusion_matrix
from sklearn.metrics import confusion_matrix,classification_report
from sklearn.metrics import balanced_accuracy_score
from sklearn.model_selection import ShuffleSplit
import random
n_folds = 5
range_N = np.arange(1, 30, 1)
p= np.arange(1,2000,100)
grid_param = {'n_neighbors': range_N, 'p': p}
cv = ShuffleSplit(n_splits=5, random_state=42)
grid_clf = GridSearchCV(KNeighborsClassifier(metric= 'minkowski'), grid_param, scoring = 'accuracy', cv=cv, verbose=3,  return_train_score=True)
grid_clf.fit(scaled_features, Y)
best_grid = grid_clf.best_params_.copy()
print("Selected settings: ", best_grid)
print("CV-score: %.2f" % grid_clf.best_score_)
KNN = grid_clf.best_estimator_
y_pred_train = KNN.predict(scaled_features)
y_pred_test = KNN.predict(X_test)
acc_train = balanced_accuracy_score(Y, y_pred_train)
acc_test = balanced_accuracy_score(Y_test, y_pred_test)
print("Train/test accuracy: %.2f/%.2f" % (acc_train, acc_test))
print(" Train Recall: {:.2f}".format(recall_score(Y, y_pred_train, average='macro')))
print("Train Precision: {:.2f}".format(precision_score(Y, y_pred_train, average='macro')))
print("Train f1_score: {:.2f}".format(f1_score(Y, y_pred_train, average='macro')))
print("Test Recall: {:.2f}".format(recall_score(Y_test, y_pred_test, average='macro')))
print("Test Precision: {:.2f}".format(precision_score(Y_test, y_pred_test, average='macro')))
print("Test f1_score: {:.2f}".format(f1_score(Y_test, y_pred_test, average='macro')))
plot_confusion_matrix(KNN, scaled_features, Y)
plot_confusion_matrix(KNN, X_test, Y_test)

"""# Scoring - Used in all predictors!"""

from sklearn.metrics import roc_curve
from sklearn.metrics import make_scorer, accuracy_score, precision_score, recall_score, f1_score
from sklearn.model_selection import cross_validate
def weird_division(n, d):
    return n / d if d else 0
def false_positive_rate(y_true, y_pred):

    # false positive
    fp = ((y_pred == 1) & (y_true == -1)).sum()
    fp=np.array(fp)

    # true negative
    tn = ((y_pred == -1) & (y_true == -1)).sum()
    tn=np.array(tn)

    # false positive rate
    return 1- weird_division(fp, fp+tn)
scoring = {
    'accuracy': make_scorer(accuracy_score),
    'f1_score': make_scorer(f1_score, average='macro'),
    'false_positive_rate': make_scorer(false_positive_rate),
    }

"""# Shuffling - KNN"""

from pandas.core.common import random_state
import statistics
from sklearn.neighbors import KNeighborsClassifier
from sklearn.model_selection import cross_val_score
from sklearn.metrics import make_scorer, accuracy_score, precision_score, recall_score, f1_score
from sklearn.model_selection import KFold
from sklearn.model_selection import cross_validate
from sklearn.metrics import plot_confusion_matrix
from sklearn.metrics import confusion_matrix,classification_report
from sklearn.metrics import balanced_accuracy_score
from sklearn.model_selection import ShuffleSplit
import random
n=100
accuracy=[]
acc_train=[]
acc_test=[]
aa=[]
cc=[]
FPR_train=[]
FPR_test=[]
for j in range(n): 
  KNN = KNeighborsClassifier(n_neighbors=6, metric= 'minkowski', p=801)
  KNN.fit(scaled_features, Y)
  cv = ShuffleSplit(n_splits=5, random_state=random.seed(1234))
  scores = cross_validate(KNN, scaled_features, Y, cv=cv, scoring=scoring)
  accuracy. append(np.mean(scores["test_accuracy"]))
  y_pred_train = KNN.predict(scaled_features)
  y_pred_test = KNN.predict(X_test)
  acc_train. append(accuracy_score(Y, y_pred_train))
  acc_test. append(accuracy_score(Y_test, y_pred_test))
  aa. append(f1_score(Y_test, y_pred_test, average='macro'))
  cc.  append(np.mean(scores["test_f1_score"]))
  tn, fp, fn, tp = confusion_matrix(list(Y), list(y_pred_train), labels=[-1, 1]).ravel()
  FPR_train. append(np.mean(scores["test_false_positive_rate"]))
  tn1, fp1, fn1, tp1 = confusion_matrix(list(Y_test), list(y_pred_test), labels=[-1, 1]).ravel()
  FPR_test. append(1-(fp1/(tn1+fp1)))
print(FPR_train)
print("Train/test accuracy: %.2f/%.2f" % (np.mean(accuracy), np.mean(acc_test)))
print("Train accuracy Standard Deviation: %.2f" % statistics.pstdev(accuracy))
print("Test accuracy Standard Deviation: %.2f" % statistics.pstdev(acc_test))
print("Train f1_score : {:.2f}".format(np.mean(cc)))
print("Test f1_score: {:.2f}".format(np.mean(aa)))
print("Train f1_score Standard Deviation: %.2f" % statistics.pstdev(cc))
print("Test f1_score Standard Deviation: %.2f" % statistics.pstdev(aa))
print("Train 1-FPR : {:.2f}".format(np.mean(FPR_train)))
print("test 1-FPR : {:.2f}".format(np.mean(FPR_test)))
print("Train 1-FPR Standard Deviation: %.2f" % statistics.pstdev(FPR_train))
print("Train 1-FPR Standard Deviation: %.2f" % statistics.pstdev(FPR_test))

"""# KNN - LC"""

from sklearn.model_selection import learning_curve
from sklearn.svm import SVC
from sklearn.model_selection import cross_val_score
from sklearn.metrics import make_scorer, accuracy_score, precision_score, recall_score, f1_score
from sklearn.model_selection import KFold
from sklearn.model_selection import cross_validate
from sklearn.metrics import plot_confusion_matrix
clf = KNeighborsClassifier(metric= 'minkowski', n_neighbors= 17, p=601)
n_folds = 5
train_sizes = np.linspace(.1, 1.0, 6)
train_sizes, train_scores, test_scores, fit_times, _ = \
learning_curve(clf, X, Y,
cv=n_folds, train_sizes=train_sizes,
scoring='accuracy',
return_times=True)
train_scores_mean = np.mean(train_scores, axis=1)
train_scores_std = np.std(train_scores, axis=1)
test_scores_mean = np.mean(test_scores, axis=1)
test_scores_std = np.std(test_scores, axis=1)
fit_times_mean = np.mean(fit_times, axis=1)
fit_times_std = np.std(fit_times, axis=1)
_, axes = plt.subplots(1, 1, figsize=(10, 10))
axes.grid()
axes.fill_between(train_sizes, train_scores_mean - train_scores_std,
train_scores_mean + train_scores_std, alpha=0.1,
color="r")
axes.fill_between(train_sizes, test_scores_mean - test_scores_std,
test_scores_mean + test_scores_std, alpha=0.1,
color="g")
axes.plot(train_sizes, train_scores_mean, 'o-', color="r",
label="Training score")
axes.plot(train_sizes, test_scores_mean, 'o-', color="g",
label="Cross-validation score")
axes.legend(prop={"size":25}, loc='upper right')
#legend.get_title().set_fontsize('6')
#axes[0].legend(loc="best")
axes.set_xlabel("Training examples", fontsize= 30)
axes.set_ylabel("Accuracy", fontsize= 30)
axes.set_title("Data sensitivity", fontsize= 30)
#print(np.mean(train_scores_std))
plt.ylim(0 , 1.01)
plt.xticks(size = 20)
plt.yticks(size = 20)
plt.show()

"""# Making the files of predictions"""

from sklearn.neighbors import KNeighborsClassifier
from sklearn.svm import SVC
from sklearn.ensemble import RandomForestClassifier
from sklearn.neural_network import MLPClassifier
from sklearn.naive_bayes import GaussianNB


KNN = KNeighborsClassifier(n_neighbors=6, metric= 'minkowski', p=801)
clf= SVC(C=0.14, kernel="linear")
RF = RandomForestClassifier(max_depth=2, n_estimators= 4)
MLP= MLPClassifier(hidden_layer_sizes= 21, activation="tanh", max_iter=5000, solver= "sgd", alpha=1)
GB = GaussianNB()

KNN2 = KNeighborsClassifier(n_neighbors=17, metric= 'minkowski', p=601)
clf2= SVC(C=0.13, kernel="linear")
RF2 = RandomForestClassifier(max_depth=2, n_estimators= 4)
MLP2= MLPClassifier(hidden_layer_sizes= 43, activation="tanh", max_iter=5000, solver= "sgd", alpha=0.1)
GB2 = GaussianNB()


KNN.fit(scaled_features, Y)
clf.fit(scaled_features, Y)
RF.fit(scaled_features, Y)
MLP.fit(scaled_features, Y)
GB.fit(scaled_features, Y)

KNN2.fit(scaled_features2, Y2)
clf2.fit(scaled_features2, Y2)
RF2.fit(scaled_features2, Y2)
MLP2.fit(scaled_features2, Y2)
GB2.fit(scaled_features2, Y2)


ynewl_KNN=KNN.predict(scaled_features)
ynewl_test_KNN=KNN.predict(X_test)
ynewl_SVM=clf.predict(scaled_features)
ynewl_test_SVM=clf.predict(X_test)
ynewl_RF=RF.predict(scaled_features)
ynewl_test_RF=RF.predict(X_test)
ynewl_MLP=MLP.predict(scaled_features)
ynewl_test_MLP=MLP.predict(X_test)
ynewl_GB=GB.predict(scaled_features)
ynewl_test_GB=GB.predict(X_test)

ynewl_KNN2=KNN2.predict(scaled_features2)
ynewl_test_KNN2=KNN2.predict(X_test2)
ynewl_SVM2=clf2.predict(scaled_features2)
ynewl_test_SVM2=clf2.predict(X_test2)
ynewl_RF2=RF2.predict(scaled_features2)
ynewl_test_RF2=RF2.predict(X_test2)
ynewl_MLP2=MLP2.predict(scaled_features2)
ynewl_test_MLP2=MLP2.predict(X_test2)
ynewl_GB2=GB2.predict(scaled_features2)
ynewl_test_GB2=GB2.predict(X_test2)

df=df.iloc[:, 0:23]
df2=df2.iloc[:, 0:23]

df['3class_KNN']= ynewl_KNN
df_test['3class_KNN']= ynewl_test_KNN
df['3class_SVM']= ynewl_SVM
df_test['3class_SVM']= ynewl_test_SVM
df['3class_RF']= ynewl_RF
df_test['3class_RF']= ynewl_test_RF
df['3class_NN']= ynewl_MLP
df_test['3class_NN']= ynewl_test_MLP
df['3class_GB']= ynewl_GB
df_test['3class_GB']= ynewl_test_GB

df2['2class_KNN']= ynewl_KNN2
df_test2['2class_KNN']= ynewl_test_KNN2
df2['2class_SVM']= ynewl_SVM2
df_test2['2class_SVM']= ynewl_test_SVM2
df2['2class_RF']= ynewl_RF2
df_test2['2class_RF']= ynewl_test_RF2
df2['2class_NN']= ynewl_MLP2
df_test2['2class_NN']= ynewl_test_MLP2
df2['2class_GB']= ynewl_GB2
df_test2['2class_GB']= ynewl_test_GB2
  #print(proteins[i], ynewl[i], Y_test[i])
print (df)
print (df_test)
print (df2)
print (df_test2)

"""# Download the files"""

from google.colab import files

df.to_csv('df.csv')
files.download('df.csv')

df_test.to_csv('df_test.csv')
files.download('df_test.csv')

df2.to_csv('df2.csv')
files.download('df2.csv')

df_test2.to_csv('df_test2.csv')
files.download('df_test2.csv')

"""# Grid_Param_SVM"""

range_C = np.logspace(-3, 1, num=1000)
print(range_C)
kernel=['rbf', 'poly', 'linear']
range_C = np.logspace(-3, 0, num=50)
print(range_C)
grid_param = {'C': range_C, 'kernel': kernel}

"""# SVM"""

from sklearn.model_selection import GridSearchCV
from sklearn.svm import SVC
from sklearn.model_selection import cross_val_score
from sklearn.metrics import make_scorer, accuracy_score, precision_score, recall_score, f1_score
from sklearn.model_selection import KFold
from sklearn.model_selection import cross_validate
from sklearn.metrics import plot_confusion_matrix
from sklearn.metrics import confusion_matrix,classification_report
from sklearn.metrics import balanced_accuracy_score
n_folds=5
range_C = np.logspace(-3, 0, num=500)
grid_param = {'C': range_C, 'kernel': kernel}
n=100
accuracy=[]
acc_train=[]
acc_test=[]
aa=[]
grid_clf = GridSearchCV(SVC(), grid_param, scoring = 'accuracy', cv=n_folds, verbose=3,  return_train_score=True)
grid_clf.fit(scaled_features, Y)
best_grid = grid_clf.best_params_.copy()
print("Selected settings: ", best_grid)
print("CV-score: %.2f" % grid_clf.best_score_)
clf = grid_clf.best_estimator_
y_pred_train = clf.predict(scaled_features)
y_pred_test = clf.predict(X_test)
acc_train = balanced_accuracy_score(Y, y_pred_train)
acc_test = balanced_accuracy_score(Y_test, y_pred_test)
print("Train/test accuracy: %.2f/%.2f" % (acc_train, acc_test))
print(" Train Recall: {:.2f}".format(recall_score(Y, y_pred_train, average='macro')))
print("Train Precision: {:.2f}".format(precision_score(Y, y_pred_train, average='macro')))
print("Train f1_score: {:.2f}".format(f1_score(Y, y_pred_train, average='macro')))
print("Test Recall: {:.2f}".format(recall_score(Y_test, y_pred_test, average='macro')))
print("Test Precision: {:.2f}".format(precision_score(Y_test, y_pred_test, average='macro')))
print("Test f1_score: {:.2f}".format(f1_score(Y_test, y_pred_test, average='macro')))
plot_confusion_matrix(clf, scaled_features, Y)
plot_confusion_matrix(clf, X_test, Y_test)

"""# Shuffling - SVM"""

from sklearn.model_selection import ShuffleSplit
import statistics
n=100
accuracy=[]
acc_train=[]
acc_test=[]
aa=[]
cc=[]
FPR_train=[]
FPR_test=[]
for j in range(n): 
  clf = SVC(C=0.14, kernel='linear')
  clf.fit(scaled_features, Y)
  y_pred_train = clf.predict(scaled_features)
  FPR = make_scorer(false_positive_rate(Y, y_pred_train))
  cv = ShuffleSplit(n_splits=5, random_state=random.seed(1234))
  scores = cross_validate(clf, scaled_features, Y, cv=cv,scoring= scoring)
  accuracy. append(np.mean(scores['test_accuracy']))
  y_pred_train = clf.predict(scaled_features)
  y_pred_test = clf.predict(X_test)
  acc_train. append(accuracy_score(Y, y_pred_train))
  acc_test. append(accuracy_score(Y_test, y_pred_test))
  aa. append(f1_score(Y_test, y_pred_test, average='macro'))
  cc. append(np.mean(scores['test_f1_score']))
  tn, fp, fn, tp = confusion_matrix(list(Y), list(y_pred_train), labels=[-1, 1]).ravel()
  FPR_train. append(np.mean(scores['test_false_positive_rate']))
  tn1, fp1, fn1, tp1 = confusion_matrix(list(Y_test), list(y_pred_test), labels=[-1, 1]).ravel()
  FPR_test. append(1-(fp1/(tn1+fp1)))
print(FPR_train)
print("Train/test accuracy: %.2f/%.2f" % (np.mean(accuracy), np.mean(acc_test)))
print("Train accuracy Standard Deviation: %.2f" % statistics.pstdev(accuracy))
print("Test accuracy Standard Deviation: %.2f" % statistics.pstdev(acc_test))
print("Train f1_score : {:.2f}".format(np.mean(cc)))
print("Test f1_score: {:.2f}".format(np.mean(aa)))
print("Train f1_score Standard Deviation: %.2f" % statistics.pstdev(cc))
print("Test f1_score Standard Deviation: %.2f" % statistics.pstdev(aa))
print("Train 1-FPR : {:.2f}".format(np.mean(FPR_train)))
print("test 1-FPR : {:.2f}".format(np.mean(FPR_test)))
print("Train 1-FPR Standard Deviation: %.2f" % statistics.pstdev(FPR_train))
print("Train 1-FPR Standard Deviation: %.2f" % statistics.pstdev(FPR_test))

"""# SVM - LC"""

from sklearn.model_selection import learning_curve
from sklearn.svm import SVC
from sklearn.model_selection import cross_val_score
from sklearn.metrics import make_scorer, accuracy_score, precision_score, recall_score, f1_score
from sklearn.model_selection import KFold
from sklearn.model_selection import cross_validate
from sklearn.metrics import plot_confusion_matrix
clf = SVC(C=0.13, kernel='linear', probability=True)
n_folds = 5
train_sizes = np.linspace(.1, 1.0, 6)
train_sizes, train_scores, test_scores, fit_times, _ = \
learning_curve(clf, scaled_features, Y,
cv=n_folds, train_sizes=train_sizes,
scoring='accuracy',
return_times=True)
train_scores_mean = np.mean(train_scores, axis=1)
train_scores_std = np.std(train_scores, axis=1)
test_scores_mean = np.mean(test_scores, axis=1)
test_scores_std = np.std(test_scores, axis=1)
fit_times_mean = np.mean(fit_times, axis=1)
fit_times_std = np.std(fit_times, axis=1)
_, axes = plt.subplots(1, 1, figsize=(10, 10))
axes.grid()
axes.fill_between(train_sizes, train_scores_mean - train_scores_std,
train_scores_mean + train_scores_std, alpha=0.1,
color="r")
axes.fill_between(train_sizes, test_scores_mean - test_scores_std,
test_scores_mean + test_scores_std, alpha=0.1,
color="g")
axes.plot(train_sizes, train_scores_mean, 'o-', color="r",
label="Training score")
axes.plot(train_sizes, test_scores_mean, 'o-', color="g",
label="Cross-validation score")
axes.legend(prop={"size":25}, loc='upper right')
#legend.get_title().set_fontsize('6')
#axes[0].legend(loc="best")
axes.set_xlabel("Training examples", fontsize= 30)
axes.set_ylabel("Accuracy", fontsize= 30)
axes.set_title("Data sensitivity", fontsize= 30)
#print(np.mean(train_scores_std))
plt.xlim(40 , 165)
plt.ylim(0 , 1)
plt.xticks(size = 20)
plt.yticks(size = 20)
plt.show()

"""# Grid_param - RF"""

range_depth = np.arange(1, 5, 1)
range_trees = np.arange(1, 5, 1)
#range_criterion = ['gini', 'entropy']
grid_param = {'n_estimators' : range_trees,'max_depth': range_depth}

"""# RF and Rf - LC"""

from sklearn.ensemble import RandomForestClassifier
from sklearn.model_selection import learning_curve
from sklearn.model_selection import GridSearchCV
from sklearn.model_selection import cross_val_score
from sklearn.metrics import make_scorer, accuracy_score, precision_score, recall_score, f1_score
from sklearn.model_selection import KFold
from sklearn.model_selection import cross_validate
from sklearn.metrics import plot_confusion_matrix
n_folds=5
grid_RF = GridSearchCV(RandomForestClassifier(), grid_param, scoring = 'accuracy', cv=n_folds)
grid_RF.fit(scaled_features, Y)
best_grid = grid_RF.best_params_.copy()
print("Selected settings: ", best_grid)
print("CV-score: %.2f" % grid_RF.best_score_)
RF = grid_RF.best_estimator_
y_pred_train = RF.predict(scaled_features)
y_pred_test = RF.predict(X_test)
acc_train = balanced_accuracy_score(Y, y_pred_train)
acc_test = balanced_accuracy_score(Y_test, y_pred_test)
print("Train/test accuracy: %.2f/%.2f" % (acc_train, acc_test))
print("Train Recall: {:.2f}".format(recall_score(Y, y_pred_train, average='micro')))
print("Train Precision: {:.2f}".format(precision_score(Y, y_pred_train, average='micro')))
print("Train f1_score: {:.2f}".format(f1_score(Y, y_pred_train, average='micro')))
print("Test Recall: {:.2f}".format(recall_score(Y_test, y_pred_test, average='micro')))
print("Test Precision: {:.2f}".format(precision_score(Y_test, y_pred_test, average='micro')))
print("Test f1_score: {:.2f}".format(f1_score(Y_test, y_pred_test, average='micro')))
plot_confusion_matrix(RF, scaled_features, Y)
plot_confusion_matrix(RF, X_test, Y_test)
from sklearn.model_selection import learning_curve
from sklearn.svm import SVC
from sklearn.model_selection import cross_val_score
from sklearn.metrics import make_scorer, accuracy_score, precision_score, recall_score, f1_score
from sklearn.model_selection import KFold
from sklearn.model_selection import cross_validate
from sklearn.metrics import plot_confusion_matrix
train_sizes = np.linspace(.1, 1.0, 6)
train_sizes, train_scores, test_scores, fit_times, _ = \
learning_curve(RF, scaled_features, Y,
cv=n_folds, train_sizes=train_sizes,
scoring='accuracy',
return_times=True)
train_scores_mean = np.mean(train_scores, axis=1)
train_scores_std = np.std(train_scores, axis=1)
test_scores_mean = np.mean(test_scores, axis=1)
test_scores_std = np.std(test_scores, axis=1)
fit_times_mean = np.mean(fit_times, axis=1)
fit_times_std = np.std(fit_times, axis=1)
_, axes = plt.subplots(1, 1, figsize=(10, 10))
axes.grid()
axes.fill_between(train_sizes, train_scores_mean - train_scores_std,
train_scores_mean + train_scores_std, alpha=0.1,
color="r")
axes.fill_between(train_sizes, test_scores_mean - test_scores_std,
test_scores_mean + test_scores_std, alpha=0.1,
color="g")
axes.plot(train_sizes, train_scores_mean, 'o-', color="r",
label="Training score")
axes.plot(train_sizes, test_scores_mean, 'o-', color="g",
label="Cross-validation score")
axes.legend(prop={"size":25}, loc='upper right')
#legend.get_title().set_fontsize('6')
#axes[0].legend(loc="best")
axes.set_xlabel("Training examples", fontsize= 30)
axes.set_ylabel("Accuracy", fontsize= 30)
axes.set_title("Data sensitivity", fontsize= 30)
plt.xlim(40 , 165)
plt.ylim(0 , 1.01)
plt.xticks(size = 20)
plt.yticks(size = 20)
#print(np.mean(train_scores_std))
plt.show()

"""# Shuffling - RF"""

import statistics
from sklearn.ensemble import RandomForestClassifier
import random
from sklearn.model_selection import cross_val_score
from sklearn.metrics import balanced_accuracy_score
from sklearn.metrics import f1_score
from sklearn.metrics import confusion_matrix
n=100
accuracy=[]
acc_train=[]
acc_test=[]
aa=[]
cc=[]
FPR_train=[]
FPR_test=[]
for j in range(n): 
  RF = RandomForestClassifier(max_depth=2, n_estimators= 4, random_state=random.seed(1234))
  RF.fit(scaled_features, Y)
  scores = cross_val_score(RF, scaled_features, Y, cv=5,scoring='accuracy')
  accuracy. append(np.mean(scores))
  y_pred_train = RF.predict(scaled_features)
  y_pred_test = RF.predict(X_test)
  acc_train. append(balanced_accuracy_score(Y, y_pred_train))
  acc_test. append(balanced_accuracy_score(Y_test, y_pred_test))
  aa. append(f1_score(Y_test, y_pred_test, average='macro'))
  cc. append(f1_score(Y, y_pred_train, average='macro'))
  tn, fp, fn, tp = confusion_matrix(list(Y), list(y_pred_train), labels=[-1, 1]).ravel()
  FPR_train. append(1-(fp/(tn+fp)))
  tn1, fp1, fn1, tp1 = confusion_matrix(list(Y_test), list(y_pred_test), labels=[-1, 1]).ravel()
  FPR_test. append(1-(fp1/(tn1+fp1)))
print(FPR_train)
print("Train/test accuracy: %.2f/%.2f" % (np.mean(accuracy), np.mean(acc_test)))
print("Train accuracy Standard Deviation: %.2f" % statistics.pstdev(accuracy))
print("Test accuracy Standard Deviation: %.2f" % statistics.pstdev(acc_test))
print("Train f1_score : {:.2f}".format(np.mean(cc)))
print("Test f1_score: {:.2f}".format(np.mean(aa)))
print("Train f1_score Standard Deviation: %.2f" % statistics.pstdev(cc))
print("Test f1_score Standard Deviation: %.2f" % statistics.pstdev(aa))
print("Train 1-FPR : {:.2f}".format(np.mean(FPR_train)))
print("test 1-FPR : {:.2f}".format(np.mean(FPR_test)))
print("Train 1-FPR Standard Deviation: %.2f" % statistics.pstdev(FPR_train))
print("Train 1-FPR Standard Deviation: %.2f" % statistics.pstdev(FPR_test))

"""# Grid_param - ANN"""

range_hidden = np.arange(2, 64, 1)
range_alpha= np.logspace(-1,0, num=10)
grid_param = [
        {
            #'activation' : ['identity', 'logistic', 'tanh', 'relu'],
            #'solver' : ['lbfgs', 'sgd', 'adam'],
            'hidden_layer_sizes': range_hidden,
            'alpha' : range_alpha
            
        }
       ]

"""# ANN and ANN-LC"""

from sklearn.model_selection import GridSearchCV
from sklearn.metrics import confusion_matrix,classification_report
from sklearn.neural_network import MLPClassifier
from sklearn.model_selection import cross_val_score
from sklearn.metrics import make_scorer, accuracy_score, precision_score, recall_score, f1_score
from sklearn.model_selection import KFold
from sklearn.model_selection import cross_validate
from sklearn.metrics import plot_confusion_matrix
from sklearn.metrics import balanced_accuracy_score
import pickle
n_folds=5
grid_MLP = GridSearchCV(MLPClassifier(activation="tanh", max_iter=5000, solver= "sgd"), grid_param, scoring = 'accuracy', cv=n_folds, verbose=3,  return_train_score=True)
grid_MLP.fit(scaled_features, Y)
best_grid = grid_MLP.best_params_.copy()
print("Selected settings: ", best_grid)
print("CV-score: %.2f" % grid_MLP.best_score_)
MLP = grid_MLP.best_estimator_
y_pred_train = MLP.predict(scaled_features)
y_pred_test = MLP.predict(X_test)
acc_train = balanced_accuracy_score(Y, y_pred_train)
acc_test = balanced_accuracy_score(Y_test, y_pred_test)
print("Train/test accuracy: %.2f/%.2f" % (acc_train, acc_test))
print("Train Recall: {:.2f}".format(recall_score(Y, y_pred_train, average='micro')))
print("Train Precision: {:.2f}".format(precision_score(Y, y_pred_train, average='micro')))
print("Train f1_score: {:.2f}".format(f1_score(Y, y_pred_train, average='micro')))
print("Test Recall: {:.2f}".format(recall_score(Y_test, y_pred_test, average='micro')))
print("Test Precision: {:.2f}".format(precision_score(Y_test, y_pred_test, average='micro')))
print("Test f1_score: {:.2f}".format(f1_score(Y_test, y_pred_test, average='micro')))
plot_confusion_matrix(MLP, scaled_features, Y)
plot_confusion_matrix(MLP, X_test, Y_test)
from sklearn.model_selection import learning_curve
from sklearn.svm import SVC
from sklearn.model_selection import cross_val_score
from sklearn.metrics import make_scorer, accuracy_score, precision_score, recall_score, f1_score
from sklearn.model_selection import KFold
from sklearn.model_selection import cross_validate
from sklearn.metrics import plot_confusion_matrix
train_sizes = np.linspace(.1, 1.0, 6)
train_sizes, train_scores, test_scores, fit_times, _ = \
learning_curve(MLP, scaled_features, Y,
cv=n_folds, train_sizes=train_sizes,
scoring='accuracy',
return_times=True)
train_scores_mean = np.mean(train_scores, axis=1)
train_scores_std = np.std(train_scores, axis=1)
test_scores_mean = np.mean(test_scores, axis=1)
test_scores_std = np.std(test_scores, axis=1)
fit_times_mean = np.mean(fit_times, axis=1)
fit_times_std = np.std(fit_times, axis=1)
_, axes = plt.subplots(1, 1, figsize=(10, 10))
axes.grid()
axes.fill_between(train_sizes, train_scores_mean - train_scores_std,
train_scores_mean + train_scores_std, alpha=0.1,
color="r")
axes.fill_between(train_sizes, test_scores_mean - test_scores_std,
test_scores_mean + test_scores_std, alpha=0.1,
color="g")
axes.plot(train_sizes, train_scores_mean, 'o-', color="r",
label="Training score")
axes.plot(train_sizes, test_scores_mean, 'o-', color="g",
label="Cross-validation score")
axes.legend(prop={"size":25}, loc='upper right')
#legend.get_title().set_fontsize('6')
#axes[0].legend(loc="best")
axes.set_xlabel("Training examples", fontsize= 30)
axes.set_ylabel("Accuracy", fontsize= 30)
axes.set_title("Data sensitivity", fontsize= 30)
plt.xlim(40 , 165)
plt.ylim(0 , 1.01)
plt.xticks(size = 20)
plt.yticks(size = 20)
#axes.legend(bbox_to_anchor=(1, 1), bbox_transform=plt.transFigure)
#print(np.mean(train_scores_std))
plt.show()

"""# Shuffling - ANN"""

import statistics
from sklearn.neural_network import MLPClassifier
from sklearn.model_selection import cross_val_score
from sklearn.metrics import make_scorer, accuracy_score, precision_score, recall_score, f1_score
from sklearn.model_selection import KFold
from sklearn.model_selection import cross_validate
from sklearn.metrics import plot_confusion_matrix
from sklearn.metrics import confusion_matrix
n=100
accuracy=[]
acc_train=[]
acc_test=[]
aa=[]
cc=[]
FPR_train=[]
FPR_test=[]
for j in range(n): 
  MLP = MLPClassifier(activation= 'tanh', solver= 'sgd', alpha=1, hidden_layer_sizes=21, max_iter=5000, random_state=random.seed(1234))
  MLP.fit(scaled_features, Y)
  scores = cross_val_score(MLP, scaled_features, Y, cv=5,scoring='accuracy')
  accuracy. append(np.mean(scores))
  y_pred_train = MLP.predict(scaled_features)
  y_pred_test = MLP.predict(X_test)
  acc_train. append(accuracy_score(Y, y_pred_train))
  acc_test. append(accuracy_score(Y_test, y_pred_test))
  aa. append(f1_score(Y_test, y_pred_test, average='weighted'))
  cc. append(f1_score(Y, y_pred_train, average='weighted'))
  tn, fp, fn, tp = confusion_matrix(list(Y), list(y_pred_train), labels=[-1, 1]).ravel()
  FPR_train. append(1-(fp/(tn+fp)))
  tn1, fp1, fn1, tp1 = confusion_matrix(list(Y_test), list(y_pred_test), labels=[-1, 1]).ravel()
  FPR_test. append(1-(fp1/(tn1+fp1)))
print(accuracy)
print("Train/test accuracy: %.4f/%.4f" % (np.mean(accuracy), np.mean(acc_test)))
print("Train accuracy Standard Deviation: %.4f" % statistics.pstdev(accuracy))
print("Test accuracy Standard Deviation: %.4f" % statistics.pstdev(acc_test))
print("Train f1_score : {:.4f}".format(np.mean(cc)))
print("Test f1_score: {:.4f}".format(np.mean(aa)))
print("Train f1_score Standard Deviation: %.4f" % statistics.pstdev(cc))
print("Test f1_score Standard Deviation: %.4f" % statistics.pstdev(aa))
print("Train 1-FPR : {:.4f}".format(np.mean(FPR_train)))
print("test 1-FPR : {:.4f}".format(np.mean(FPR_test)))
print("Train 1-FPR Standard Deviation: %.4f" % statistics.pstdev(FPR_train))
print("Train 1-FPR Standard Deviation: %.4f" % statistics.pstdev(FPR_test))

grouped = df.groupby(df['Pocket Type'])
buried = grouped.get_group(-1)
#print(buried.count())
surface = grouped.get_group(1)
#print(surface.count())
buriedX= buried.iloc[ :, 1: 22]
scaled_buried = StandardScaler().fit_transform(buriedX)
surfaceX= surface.iloc[ :, 1: 22]
scaled_surface = StandardScaler().fit_transform(surfaceX)

"""# Shuffling - GaussianNB"""

n=100
accuracy=[]
acc_train=[]
acc_test=[]
aa=[]
cc=[]
FPR_train=[]
FPR_test=[]
for j in range(n): 
  GB = GaussianNB()
  GB.fit(scaled_features, Y)
  cv = ShuffleSplit(n_splits=5, random_state=random.seed(1234))
  scores = cross_validate(GB, scaled_features, Y, cv=cv, scoring=scoring)
  accuracy. append(np.mean(scores["test_accuracy"]))  
  y_pred_train = GB.predict(scaled_features)
  y_pred_test = GB.predict(X_test)
  acc_train. append(accuracy_score(Y, y_pred_train))
  acc_test. append(accuracy_score(Y_test, y_pred_test))
  aa. append(f1_score(Y_test, y_pred_test, average='macro'))
  cc. append(np.mean(scores['test_f1_score']))
  tn, fp, fn, tp = confusion_matrix(list(Y), list(y_pred_train), labels=[-1, 1]).ravel()
  FPR_train. append(np.mean(scores['test_false_positive_rate']))
  tn1, fp1, fn1, tp1 = confusion_matrix(list(Y_test), list(y_pred_test), labels=[-1, 1]).ravel()
  FPR_test. append(1-(fp1/(tn1+fp1)))
print(accuracy)
print("Train/test accuracy: %.2f/%.2f" % (np.mean(accuracy), np.mean(acc_test)))
print("Train accuracy Standard Deviation: %.2f" % statistics.pstdev(accuracy))
print("Test accuracy Standard Deviation: %.2f" % statistics.pstdev(acc_test))
print("Train f1_score : {:.2f}".format(np.mean(cc)))
print("Test f1_score: {:.2f}".format(np.mean(aa)))
print("Train f1_score Standard Deviation: %.2f" % statistics.pstdev(cc))
print("Test f1_score Standard Deviation: %.2f" % statistics.pstdev(aa))
print("Train 1-FPR : {:.2f}".format(np.mean(FPR_train)))
print("test 1-FPR : {:.2f}".format(np.mean(FPR_test)))
print("Train 1-FPR Standard Deviation: %.2f" % statistics.pstdev(FPR_train))
print("Train 1-FPR Standard Deviation: %.2f" % statistics.pstdev(FPR_test))
```
